# The ubiquitin protease Ubp10 suppresses the formation of translocations at Cdc13 binding sites

**DOI:** 10.1101/2025.06.27.661970

**Authors:** David I. Gonzalez, Allison R. Westerbeek, Esther A. Epum, Katherine L. Friedman

## Abstract

Double strand breaks (DSBs) pose a significant threat to chromosome stability and, if left unrepaired, can result in chromosome rearrangements. Canonical DNA repair pathways mitigate these risks. However, if these repair mechanisms fail to repair the DSB, alternative repair pathways, such as break-induced replication (BIR), single-strand annealing (SSA), and *de novo* telomere addition (*dn*TA), can be utilized. Yeast subtelomeric regions are hotspots of recombination, while interstitial telomere-like sites can promote *dn*TA. In yeast, *dn*TA sites, termed SiRTAs (Sites of Repair-associated Telomere Addition), require Cdc13 association. We identified the ubiquitin protease Ubp10 as a positive regulator of *dn*TA at SiRTAs. Loss of *UBP10* reduces *dn*TA frequency but increases the frequency of other chromosomal rearrangements at SiRTAs. SiRTAs utilize the repetitive subtelomeric regions of donor chromosomes to facilitate rearrangements, with a fraction occurring independently of *RAD51* and requiring Sir4 and Sir2 components of the SIR complex. Association of Cdc13 with the SiRTA is necessary and sufficient to stimulate translocations in the absence of *UBP10.* This study highlights the diversity of DNA repair mechanisms at SiRTAs, advancing our understanding of telomere maintenance and chromosomal rearrangement formation.

**GRAPHICAL ABSTRACT:** 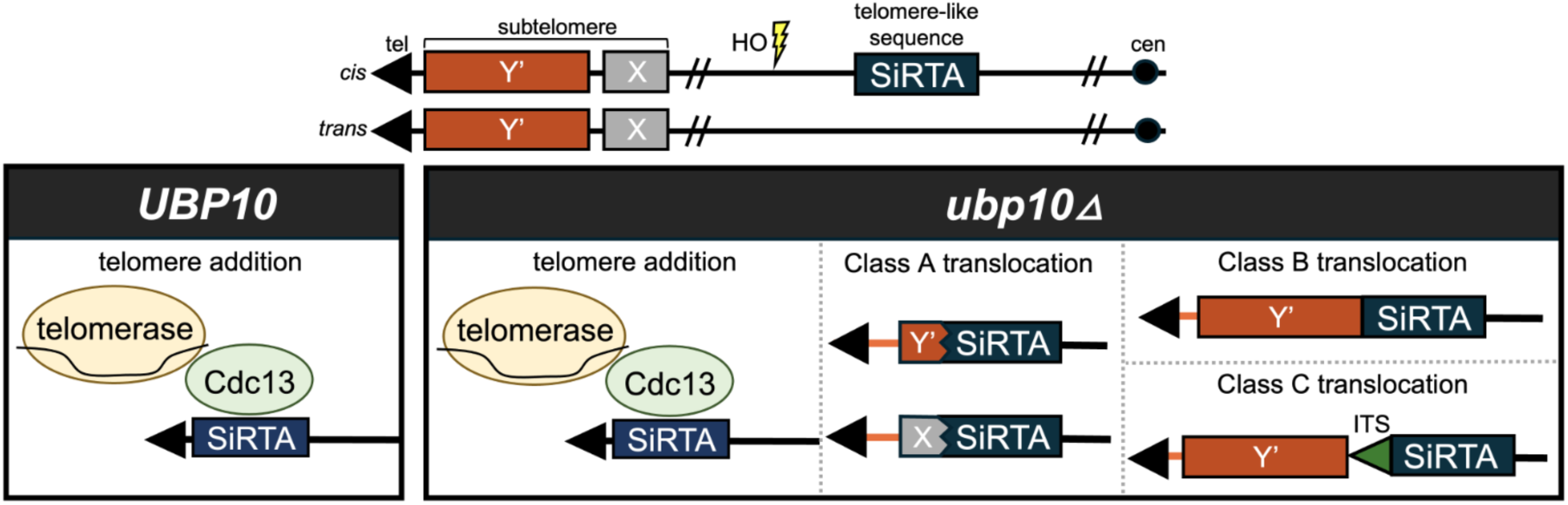

## INTRODUCTION

The maintenance of genomic integrity is crucial for human health, as alterations in the genome can lead to cancer or inherited genetic disorders. Eukaryotic chromosomes terminate in specialized nucleoprotein structures called telomeres, which play an essential role in maintaining genomic stability and integrity (1). Telomeres distinguish natural chromosome ends from processed DNA double strand breaks (DSBs), ensuring proper cellular responses to DNA damage (2). Telomeres also serve as primers for elongation by the ribonucleoprotein enzyme complex telomerase, counteracting the sequence loss inherent to each replication cycle (3). Telomerase, a reverse transcriptase, utilizes an intrinsic RNA component as a template to extend the TG-rich sequence (TG_1-3_ in yeast) of the 3’ single-stranded DNA overhang at the chromosome terminus (1, 2, 4). Subsequently, the CA-rich strand is synthesized by the lagging strand polymerase machinery (5). In budding yeast, the single-stranded DNA binding protein Cdc13 recruits and coordinates the actions of both telomerase and the lagging strand machinery (2, 6, 7). Sequence-specific recognition of the telomeric 3’ overhang by Cdc13 is required for telomere integrity and replication.

DSBs represent one of the most severe threats to genomic stability. Unrepaired DSBs can lead to mutations, chromosomal rearrangements, or cell death. Cells employ canonical DNA repair pathways to address these breaks, including homologous recombination (HR), and non-homologous end-joining (NHEJ) (8). NHEJ directly ligates the broken DNA with minimal processing, while HR utilizes a homologous template for accurate repair. For HR to proceed, the 5’-ends at a DSB must be resected to generate 3’ single-stranded DNA overhangs, which are subsequently coated by replication protein A (RPA) (9, 10). Displacement of RPA by Rad52 facilitates the assembly of the Rad51 nucleoprotein filament which conducts homology search and strand invasion into a homologous donor sequence (11, 12). While HR and NHEJ represent the primary repair mechanisms for DSB repair, cells can also invoke alternative pathways including single-strand annealing (SSA), microhomology-mediated end joining (MMEJ), break-induced replication (BIR), or *de novo* telomere addition (*dn*TA)(13–15). Among these, BIR is particularly noteworthy for its ability to repair one-ended DSBs and collapsed replication forks. Although BIR typically depends on Rad51, a rare Rad51-independent variant of BIR remains insufficiently characterized (13, 16).

BIR is prominently observed in cells utilizing the alternative lengthening of telomere (ALT) pathway. In ALT cells, telomere elongation occurs independently of telomerase through mechanisms involving BIR, often utilizing subtelomeric regions as recombination substrates (17, 18). The subtelomeric regions of yeast chromosomes are comprised of repetitive elements that facilitate recombination. All yeast subtelomeres contain an X element, followed on a subset of chromosome ends by one or more distal Y’ elements. In some cases, these repetitive elements are separated by interstitial telomeric sequences (ITSs) consisting of ∼80-150 b of TG_1-3_ (2). Significant progress has been made in understanding the types of events and genetic dependencies that support yeast ALT, but it remains a difficult phenomenon to study due to the ongoing instability generated in cells lacking telomerase (17–19).

In yeast, ITSs that perfectly match the telomeric repeat are found only within the subtelomeric sequences. However, telomere-like (TG-rich) sequences found throughout the genome can stimulate *dn*TA when exposed in single-stranded DNA following a DSB (20). Such sites of *dn*TA were first identified through studies of spontaneous rearrangements on the left arm of chromosome V (21–23). Subsequently, additional sites have been identified both experimentally and computationally (24, 25). Collectively, these sequences are named <u>Si</u>tes of <u>R</u>epair-associated <u>T</u>elomere <u>A</u>ddition (SiRTAs) for their ability to stimulate *dn*TA even when the initiating break occurs several kilobases distal to the site. Sequences predicted to function as SiRTAs are more common than expected by chance, with a particularly strong enrichment in subtelomeric regions where they may facilitate formation of a new ectopic telomere following a catastrophic telomere loss (25).

SiRTAs are bipartite: telomere addition predominantly occurs within a “Core” sequence, while a “Stim” sequence, located 5’ to the Core on the TG-rich strand, promotes *dn*TA. Multiple lines of evidence suggest that Stim sequences function through the recruitment of Cdc13, including direct binding assays *in vitro* (24). SiRTA function requires 5’ end resection following a DSB to reveal Cdc13-binding sites in single-stranded DNA, congruent with the observation that Cdc13 binding is detected at a SiRTA by chromatin immunoprecipitation only following induction of a break (24–26). In addition, artificial recruitment of Cdc13 to a modified, inactive Stim sequence rescues *dn*TA (24). Together, these experiments highlight Cdc13 as a key mediator of *dn*TA at SiRTAs. However, the potential involvement of Cdc13 in other DNA repair pathways remains insufficiently characterized, raising intriguing questions about its functional contributions to other DNA repair processes when bound to interstitial sites.

In an unpublished genetic screen designed to identify factors that stimulate *dn*TA at SiRTAs, we identified the ubiquitin protease, Ubp10. Loss of *UBP10* reduces *dn*TA frequency while concurrently increasing the frequency of other damage tolerance events at SiRTAs of various efficacies. We provide evidence that these effects are specific to SiRTAs and that Cdc13 is both necessary and sufficient (at least in some chromosome contexts) to promote translocations and/or large deletions in the absence of *UBP10.* When Ubp10 is lacking, SiRTAs utilize the repetitive subtelomeric regions of donor chromosomes to facilitate repair. Notably, some rearrangements at the SiRTA are independent of Rad51 and require Sir4 and Sir2 of the silent information regulator (SIR) complex, underscoring the diversity and complexity of DNA repair processes at these sites. This work reveals novel insights into the regulation of DNA repair mechanisms at interstitial Cdc13-bindings sites and their implications for genomic stability.

## MATERIALS AND METHODS

### Yeast strains and plasmids

Strains were constructed in the S288C background as described (24–27) and are listed in Supplement Table 1. Gene deletions were accomplished by one-step gene replacement using a selectable marker and verified by PCR and sequencing (28). Primer sequences are available in Supplement Table 2.

Strain YKF2516 was created as follows. *URA3* was amplified from pRS305 (29) and integrated on the centromere-proximal boundary of SiRTA 6L-22 using primers 6URA3F and 6URA3R. An amplicon containing the Tel11 sequence (5’-GTGTGGGTGTG-3’) and flanking DNA was generated using overlap extension PCR with primers 6L22PreSiRTA For and 6L221Cdc13BS Rev (fragment I) and 6L221Cdc13BS For and 6L22PostSiRTA Rev (fragment II). Fragments I and II were extended by mutually primed synthesis using 6L-22 PreSiRTA For and 6L22 PostSiRTA Rev. The resulting product was transformed into 6L-22::*URA3* (YKF2509), cells were allowed to recover on rich medium for 24 to 48 h, then replica plated to medium containing 5-fluoroorotic acid (5-FOA). 5-FOA resistant (FOA^R^) isolates were screened by PCR and sequenced to confirm integration of the Tel11 sequence. Strain YKF2529 was made using the same technique to replace the Stim of SiRTA 9L-44 with two tandem copies of the Gal4 upstream activating sequence (2x Gal4 UAS).

Plasmid pRS314-*UBP10* was created by inserting a fragment encompassing the *UBP10* open reading frame (Chr. XIV, nucleotides 288968 to 292101) into pRS314 using BamHI and ApaI. Plasmid pRS314-*ubp10^C371S^* was created by generating an amplicon that included the desired mutation using primers C371S For and C371S Rev. The amplicon was inserted into pRS314 using BamHI and ApaI and sequenced to confirm insertion of the desired mutation.

### Inducible HO Cleavage assay

The HO cleavage assay was performed as described (24–27, 30). Briefly, cells were grown in synthetic complete media lacking uracil (SC-Ura) containing 2% raffinose to an OD_600_ of 0.6-0.8. 10-30 μL aliquots were plated on media containing yeast extract peptone medium containing 2% galactose (YEPG). Serially diluted cells were plated on rich medium containing 2% glucose (YEPD) to determine total viable cell count. For experiments requiring plasmid selection (pRS314), cells were plated on synthetic medium lacking leucine and containing either 2% glucose or 2% galactose. Plates were incubated at 30°C for three days. Surviving colonies were counted and ∼250 galactose-resistant (Gal^R^) colonies were patched onto plates containing 5-FOA medium to isolate GCR events (Gal^R^ 5-FOA^R^ colonies). Additional Gal^R^ 5-FOA^R^ colonies were identified by replica plating when necessary. The overall GCR frequency was calculated by multiplying the frequency of Gal^R^ colonies by the fraction of Gal^R^ colonies that survived on media containing 5-FOA.

### Pooled telomere sequencing (PT-seq)

Thirty Gal^R^ 5-FOA^R^ colonies, isolated as described above, were separately inoculated in 200 μL of liquid YEPD in a 96-well culture plate and incubated overnight at 30°C. Equal volumes (at least 40 μL) of each culture were pooled and genomic DNA was extracted using the YeaStar genomic DNA kit (Zymo Research).

Libraries were prepared using 50 ng of genomic DNA and modified protocol using the Twist Library Preparation kit (Twist Bioscience 106, 543) as described in (25). DNA samples were randomly primed with 5’ barcoded adapters. Samples were pooled, captured on streptavidin-coated magnetic beads, and washed to remove excess reactants. A second 5’ adapter tailed primer with a strand-displacing polymerase was used to convert the captured templates into dual adapter libraries. Beads were washed to remove excess reactants. Four cycles of PCR were performed to amplify the library and incorporate the plate barcode in the index read position. Libraries were sequenced using the NovaSeq 6000 with 150 bp paired end reads targeting 13 to 15 million reads per sample. Real Time Analysis software (version 2.4.11; Illumina) was used for base calling and data quality control was completed using MultiQC v1.7. Library adapter sequences were removed using the Trimmomatic tool (Galaxy Version 0.38.0, (31) Using Bowtie 2 sensitive local setting (Galaxy Version 2.5.0+galaxy0), reads were aligned to the site at which the HO cut site was integrated to the last essential gene of each respective chromosome arm, referred to as the “repair region” (32, 33). Using Bowtie 2 very fast setting (Galaxy Version 2.5.0+galaxy0), reads were realigned to the *Saccharomyces cerevisiae* S288C (sacCer3) reference genome and BAM files were used for subsequent analyses (34).

### Identification, characterization, and mapping of chromosomal rearrangement breakpoints

BAM files were uploaded to the Integrative Genomic Viewer to allow visualization of reads that aligned from the HO cleavage site (HOcs) to the last essential gene of the respective chromosome arm (the “repair region”) and were arranged by mapping quality (34). GCR split reads (reads that map on one end to the repair region but also contain nucleotides aligning elsewhere in the reference genome) were classified as deletions, translocations, or telomere addition events as follows. Deletions: Split reads contain nucleotides unambiguously aligning in unique sequence located telomere proximal to the site of the *URA3* marker on the same chromosome as the HOcs. Translocations: Split reads contain nucleotides aligning to one or more genomic locations, excluding those defined as deletions. Split reads that could not unambiguously be classified as deletions were classified as translocations. Telomere addition: Split reads contain TG_1-3_ or C_1-3_A nucleotides (see examples in Supplement Figure 1A). The location of a GCR split read is defined as the genome coordinate at which the first nucleotide of the read diverges from the reference genome. The frequency of GCR split reads at a given location and/or of a given type was quantified by dividing the number of GCR split reads meeting those criteria by the total number of GCR split reads mapping from the HOcs to the last essential gene. Precise chromosome coordinates for each region analyzed are given in Figure 1 and Supplement Table 3. Previously published Nanopore sequencing data for the strains used in this study showed that the left arms of chromosomes IX, V, VI, and XIV closely resemble the S288C reference genome (26).

**Figure 1.**
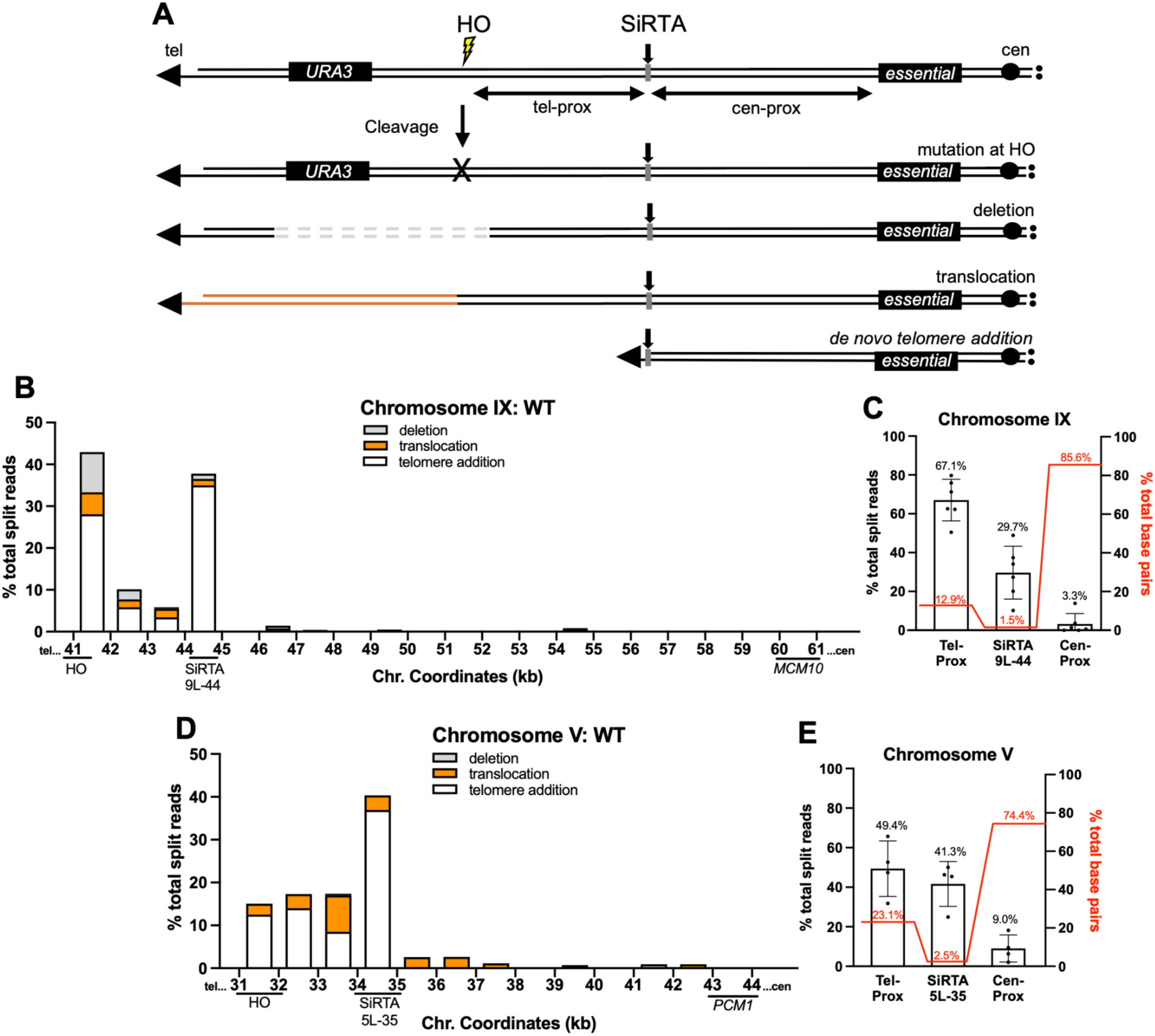
Gross chromosomal rearrangement (GCR) location and frequency are determined by mapping and quantifying split reads proximal to a site of induced cleavage. (**A**) Schematic diagram of the HO cleavage assay system. Cells that acquire resistance to galactose (used to induce expression of the HO endonuclease) and 5-FOA are assumed to have undergone a GCR event such as a deletion, translocation, or telomere addition. (**B**) Percentages of total split reads that map to each 1 kb interval within the repair region (HO cleavage site to *MCM10*, most distal essential gene on chromosome IX) are shown. Split reads were classified as deletions (gray), translocations (orange), or telomere additions (white) as described in Methods. The X axis is the distance (in kilobases) from the left end of the chromosome. Data are compiled from sequencing of 180 independent GCR events, analyzed in six pools of 30 clones each as described in Methods. (**C**) The percentages of total split reads that map telomere-proximal to SiRTA 9L-44 (41500–44100), at SiRTA 9L-44 (44100–44400), and centromere-proximal to SiRTA 9L-44 (44400–61600) are shown (left Y axis). Each data point represents a pooled sample of 30 clones that underwent a GCR event. Percentages of base pairs within the three specified regions are shown (red; right Y axis). Percentages represent the fraction of base pairs in each region relative to the total base pairs across the entire repair region. (**D**) As in (B), with cleavage targeted to chromosome V. Data are compiled from sequencing of 120 independent GCR events, analyzed in four pools. *PCM1* is the most distal essential gene. (**E**) As in (C), with split reads quantified telomere-proximal to SiRTA 5L-35 (31900–34700), at SiRTA 5L-35 (34700–35000), and centromere-proximal to SiRTA 5L-35 (35000–44000). Error bars denote Standard Deviation (SD).

Given an average number of ∼127 split reads per experiment representing 30 total GCR events, a Poisson distribution was generated to predict the number of reads expected per GCR event (Supplement Figure 1B). Using a significance level of α=0.05, two GCR split reads represented the threshold for detecting a statistically significant GCR event. Any apparent rearrangements that were represented by only one sequence read were eliminated from quantification.

## RESULTS AND DISCUSSION

### Mapping and quantification of rearrangement breakpoints following a DSB

In this work, GCR formation is monitored in haploid cells using an established assay in which a single recognition site for the HO endonuclease is integrated distal to the last essential gene on a chromosome of interest (Figure 1A). The HO endonuclease is placed under control of a galactose-inducible promoter. Cells that survive persistent nuclease expression repair the break in a way that prevents further cleavage, either through point mutations or indels within the HO cleavage site (HOcs) or through GCR events (large deletions, translocations, or telomere additions) that completely eliminate the cleavage site. By selecting for loss of a *URA3* marker integrated distal to the HOcs, we identify those clones in which a GCR event occurred between the HOcs and the most distal essential gene on that chromosome arm, here referred to as the “repair region” (Figure 1A). In previous work, we used this assay to demonstrate that SiRTAs are hot spots for telomere addition, even when located at least 3 kb proximal to the cleavage site (24–26, 35).

Previously, we validated a pooled telomere sequencing (PT-seq) approach to quantify the frequency of telomere addition events at the SiRTA (25). In brief, genomic DNA is isolated from a pool of 30 clones that survived nuclease induction on galactose, accompanied by loss of *URA3* expression (Gal^R^ 5-FOA^R^). This pooled DNA is subjected to deep, short-read (150 bp) sequencing. Among sequence reads aligning to the SiRTA, those showing evidence of a telomere addition (TG_1-3_ or C_1-3_A sequence) are counted. Read depth is normalized between experiments using read count at the most distal essential gene. This method yields results that correlate strongly with data obtained by PCR-based mapping (*r^2^=*0.97), allowing accurate estimation of the fraction of surviving cells undergoing telomere addition at the SiRTA. Our original analyses did not attempt to detect or quantify other types of rearrangements either at the SiRTA or elsewhere within the region from the HOcs to the last essential gene.

To comprehensively identify rearrangement events, we modified the analysis by incorporating the Integrative Genomic Viewer (IGV) software. Using the *S. cerevisiae* S288C reference genome (sacCer3), reads are aligned to the “repair region” extending from the site at which the HOcs was integrated to the distal boundary of the last essential gene (representing the most extensive chromosome loss event compatible with cell viability) (34). Aligned reads are arranged by mapping quality; reads with low mapping quality are predominantly split reads where the proximal portion of the read aligns to the region of interest and the distal portion aligns elsewhere in the reference genome. Split reads containing TG_1-3_ or C_1-3_A nucleotides are classified as telomere addition events (Supplementary Figure 1A). Split reads containing nucleotides unambiguously mapping to single-copy sequences distal to the *URA3* marker on the same chromosome arm are classified as deletions. Split reads containing nucleotides unambiguously mapping to locations on other chromosomes are classified as translocations. In some cases, the split read contains sequences that map to multiple locations (typically in subtelomeric repeats). These events are classified as translocations and discussed in more detail below. Because the 30 clones included in the pool have all lost the end of the cleaved chromosome (as defined by loss of *URA3*), we use the percentage of total split reads at a particular nucleotide or region as a proxy for the percentage of GCR events at that location. This approach allows us to determine the nature and frequency of chromosomal rearrangements and to map the location of each event with nucleotide precision (Figure 1B, 1D).

We applied this methodology to comprehensively map GCR events in the vicinity of two sequences previously shown to function as SiRTAs. SiRTA 9L-44 (∼44.0 kb from the endogenous telomere on the left arm of chromosome IX) lies 2.7 kb upstream of the HOcs and 17.2 kb distal to the last essential gene, *MCM10*. SiRTA 5L-35 (∼35 kb from the endogenous telomere on the left arm of chromosome V) lies 2.8 kb upstream of the HOcs and 9.0 kb from the last essential gene, *PCM1*. The “repair region” includes ∼19.9 kb on chromosome IX (HOcs to *MCM10)* and ∼12.1 kb on chromosome V (HOcs to *PCM1*). On chromosome IX, we combined data from six experiments representing a total of 180 independent GCR events. Sequence reads were aligned to the repair region and all split reads were identified and quantified. SiRTA 9L-44 comprises 1.5% of the entire repair region, yet 29.7% of total split reads map to the SiRTA (Figure 1C). Of the split reads mapping precisely to SiRTA 9L-44, 94.5% contain telomeric repeats, while the remaining 6.5% contain nucleotides indicative of a translocation/deletion (Supplement Figure 2A).

Similar results were obtained for SiRTA 5L-35. Combining results from four independent experiments (120 GCR events), 41.3% of total split reads map to the SiRTA itself (Figure 1E). SiRTA 5L-35 comprises 2.5% of the repair region on chromosome V. All split reads that map to SiRTA 5L-35 contain telomeric repeats (Supplement Figure 2B). We previously found that deletion of *RAD52* does not alter the frequency of telomere addition at SiRTAs 9L-44 and 5L-35, consistent with acquisition of telomeric sequences by telomerase rather than by recombination with telomeric or sub-telomeric sequences (an event expected to require *RAD52*) (26). Therefore, at SiRTAs 9L-44 and 5L-35, we make the simplifying assumption that all events showing acquisition of G_1-3_T sequences represent addition of a new telomere by telomerase (i.e. they are *dn*TA events).

This methodology is robust and reproducible. At SiRTA 9L-44, calculating the frequency of GCR events at the SiRTA using the percentage of total split reads that map to that sequence gives values statistically indistinguishable from values obtained using methods that normalize to total read depth or that quantify individual events by PCR (Supplement Figure 2C). We previously observed that deletion of *RAD51* increases the GCR events centromere-proximal to SiRTA 9L-44(26). Using this approach, we observed statistically indistinguishable values to those previously reported, not only at the SiRTA, but across the entire repair region (Supplement Figure 2D). Given this congruence, we use the terms “fraction of split reads” and “fraction of GCR events” interchangeably throughout this work. Importantly, GCR split reads are exclusively observed on the chromosome arm where the break is induced. Analysis of the same region on chromosome IX when the cleavage is induced on chromosome V shows no evidence of GCR split reads suggesting that GCR split reads are a result of cleavage (Supplement Table 4).

### The deubiquitinase activity of Upb10 suppresses translocations at SiRTAs

In an unpublished screen to identify *trans-*acting factors that promote *de novo* telomere addition, we identified the ubiquitin protease, Ubp10. The consequence of deleting *UBP10* was initially measured at SiRTA 9L-44. Deletion of *UBP10* does not change the fraction of total GCR events at the SiRTA, but there is a dramatic shift from *dn*TA to translocations, with translocations accounting for nearly half of the GCR events at the SiRTA (Figure 2A, Supplement Figure 3A-C). Expressing *UBP10* from its endogenous promoter on a low-copy number, centromere-containing plasmid in the *ubp10*Δ strain restores *dn*TA and eliminates translocations (Figure 2B). In contrast, expression of the catalytically dead *ubp10^C371S^* allele fails to complement the mutant defect (Figure 2B), suggesting that suppression of translocations requires the deubiquitinase activity of Ubp10.

**Figure 2.**
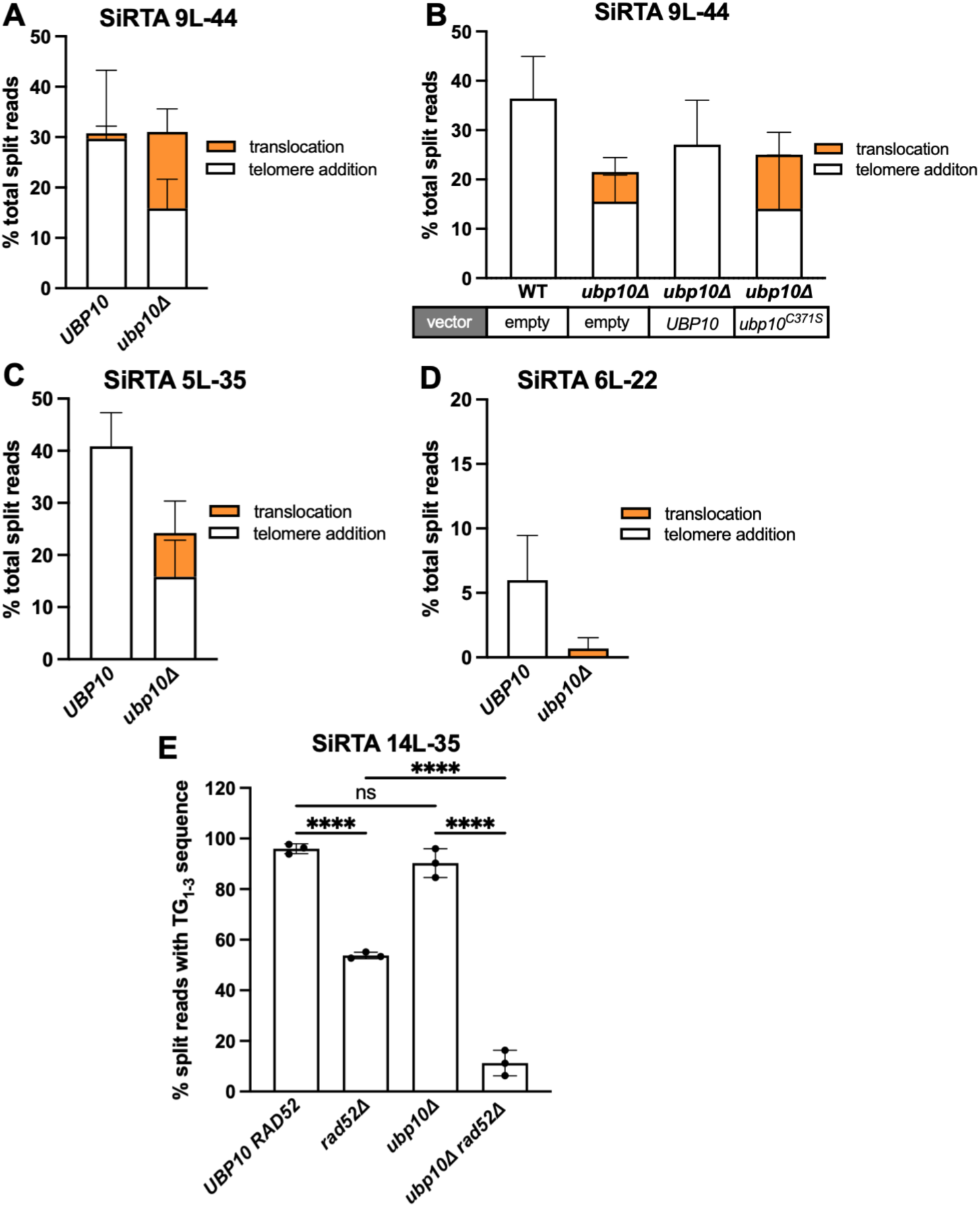
Loss of *UBP10* increases the frequency of translocations at SiRTAs. Percentages of total split reads mapping to the indicated SiRTA are shown for *UBP10* and *ubp10*Δ strains. Chromosome coordinates for each SiRTA are given in Supplement Table 3. Split reads were classified as translocations (orange) or telomere additions (white) as described in Methods. No deletions were observed. (**A**) Data for SiRTA 9L-44 are from 6 pools of 30 GCR events each (*UBP10* samples are the same as those described in Figure 1B, C). (**B**) *UBP10* and *upb10*Δ strains were transformed with either empty vector or a plasmid expressing a WT or catalytically dead variant of *UBP10* as indicated. Data are from 3 pools of 30 GCR events for each strain. (**C**) Data for SiRTA 5L-35 are from 4 pools of 30 GCR events each (*UBP10* samples are the same as those described in Figure 1D, E). (**D**) Data for SiRTA 6L-22 are from 3 pools of 30 GCR events each (30). (**E**) Percentages of total split reads mapping to SiRTA 14L-35 are shown for the indicated strains. Data are from 3 pools of 30 GCR events each. \*\*\*\**p*<0.0001, ANOVA with Tukey’s multiple comparison test. Error bars denote SD.

To determine the generalizability of this effect, we examined the impact of deleting *UBP10* at three additional SiRTAs that differ markedly in the frequency of *dnTA*. At SiRTA 5L-35, the fraction of GCR events mapping to the SiRTA decreases upon *UBP10* deletion (Figure 2C). Translocations at SiRTA 5L-35 increase (from none in the *UBP10* strain to 25.5% in the *ubp10*Δ background), but the effect is less pronounced than at SiRTA 9L-44 (compare Figure 2C to Figure 2A; Supplement Figure 3D-F). SiRTA 6L-22 (left arm of chromosome VI) stimulates frequencies of *de novo* telomere addition that marginally meet the threshold to be considered a functional SiRTA [defined as 6.6% (25, 30)]. Upon deletion of *UBP10*, telomere addition events are eliminated (Figure 2D; Supplement Figure 3G). Translocation split reads were observed in only one of three experiments at a frequency suggesting a single translocation event of 90 GCR clones analyzed (Figure 2D; Supplement Figure 3H), indicating that SiRTA 6L-22 only weakly stimulates any type of GCR event.

At the other end of the spectrum, we examined a sequence found ∼35 kb from the endogenous telomere on the left arm of chromosome XIV (SiRTA 14L-35) that stimulates high levels of telomere addition (∼95% of all GCR events) (Figure 2E). We detected no apparent difference in the frequency of telomere addition events and no translocations upon deletion of *UBP10* (Figure 2E). However, because the sequence of SiRTA 14L-35 bears high similarity to a telomeric repeat, we suspected that at least a fraction of telomere addition events assumed to arise through telomerase action might instead be translocations to subtelomeric or telomeric regions. Consistent with this hypothesis, we observe a significant (1.8-fold) decrease in the frequency of split reads at the SiRTA when *RAD52* is deleted (Figure 2E). We infer that, in contrast to our prior observations at SiRTAs 5L-35 and 9L-44 (26), nearly half of the telomere addition events at SiRTA 14L-35 occur through recombination with existing telomeric repeats. In the *ubp10*Δ background, deletion of *RAD52* causes an even more pronounced (6.9-fold) reduction in telomere addition (Figure 2E). Because the fraction of *RAD52*-dependent events (i.e. translocations) increases upon deletion of *UBP10*, we conclude that Ubp10 suppresses telomeric translocations at SiRTA 14L-35. Taken together, these results suggest that the deubiquitylation of one or more substrates by Ubp10 promotes *dn*TA by telomerase and represses formation of translocations at SiRTAs.

### Suppression of genome rearrangements by *UBP10* is most pronounced at SiRTAs

To determine whether the effects of *UBP10* deletion are specific to SiRTAs, we analyzed the types and locations of GCR events within and distal to SiRTA 9L-44 (accounting for 96.8% events arising after cleavage on chromosome IX; Figure 1C) in the presence and absence of *UBP10* (compare Figures 3A and 3B). The overall pattern of events is similar in the two strains, with no evidence of new rearrangement hotspots upon deletion of *UBP10*. This observation holds true for the region between the HOcs and SiRTA 5L-35 on chromosome V (Supplement Figures 4A and 4B), suggesting that Ubp10 affects repair pathway choice at sequences already prone to mutational repair.

**Figure 3.**
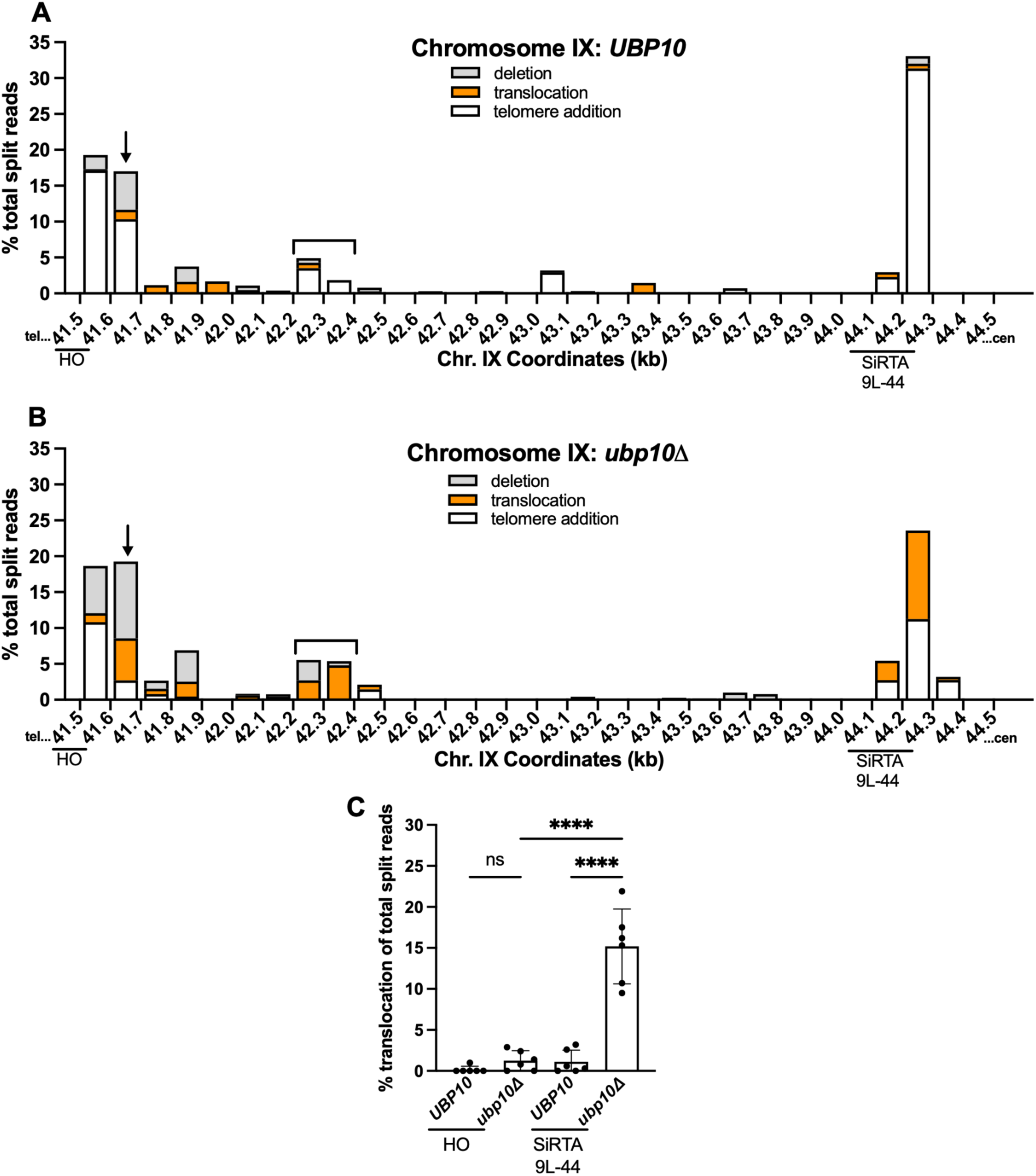
The effects of Ubp10 are most pronounced at SiRTAs. (**A-B**) Percentages of total split reads that map to each 100-bp interval between the HO cleavage site and SiRTA 9L-44 are shown. Split reads were classified as deletions (gray), translocations (orange), or telomere additions (white) as described in Methods. The X axis is the distance (in kilobases) from the left end of the chromosome. Data for *UBP10* and *ubp10*Δ strains are the same as described in Figure 2A. The arrow and brackets indicate regions where *dn*TA decreases and translocations/deletions increase upon loss of *UBP10* (see text and Supplement Figure 5). (**C**) Translocation split reads that map within 100 bp telomere proximal to the HOcs (41.5-41.6 kb) or to SiRTA 9L-44 are quantified. Percentages are calculated relative to total split reads mapping to the repair region, from the HOcs to the last essential gene. Each data point corresponds to one of the six pools of 30 GCR events as described in Figure 2A. \*\*\*\**p*<0.0001, ANOVA with Tukey’s multiple comparisons test.

**Figure 4.**
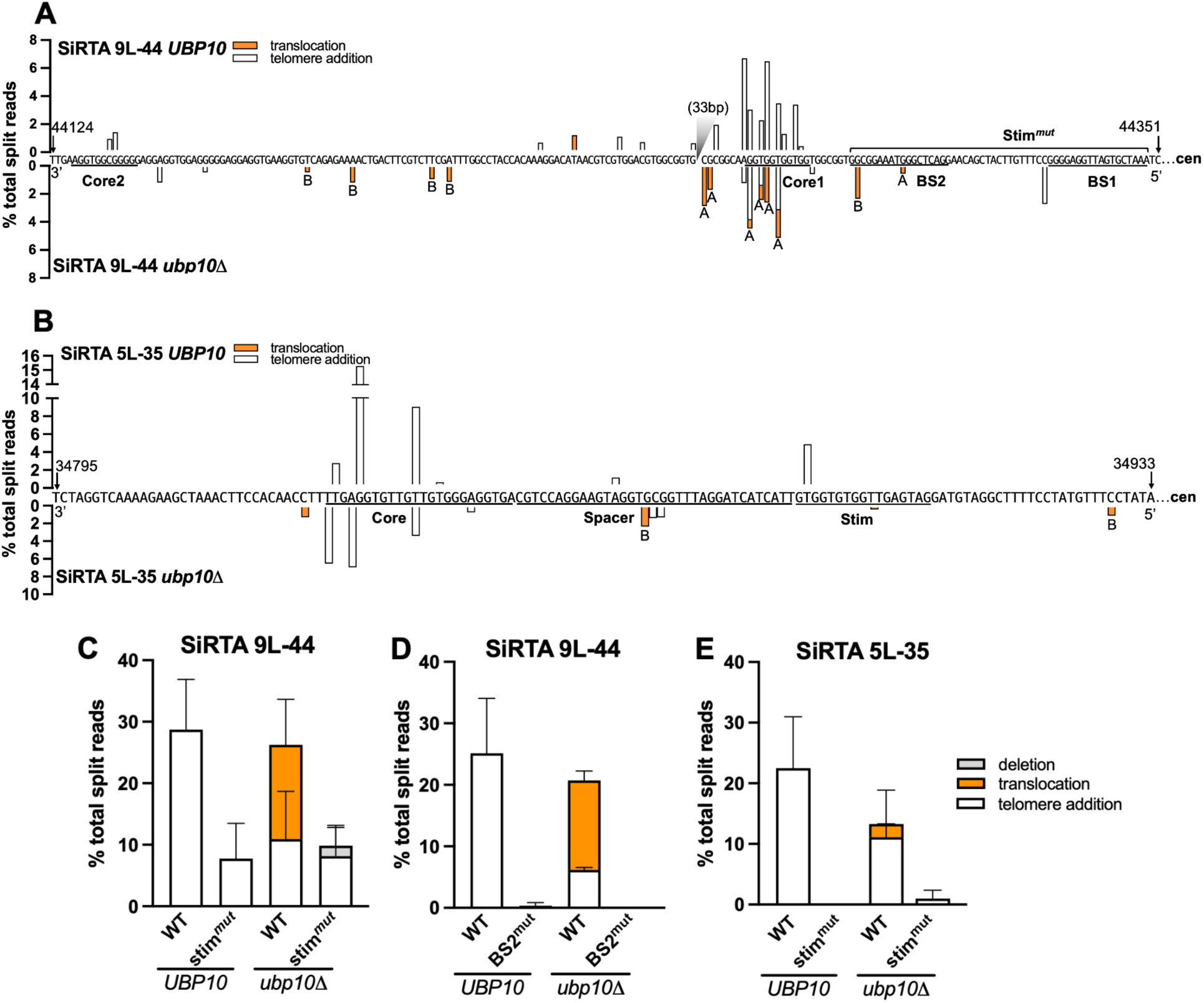
The SiRTA Stim sequence is required for translocations in the absence of *UBP10.* (**A, B**) Nucleotide-resolution maps showing sites of telomere additions (white) or translocations (orange) at 9L-44 (A) or 5L-35 (B) using the data summarized in Figure 2A and 2B. Sequences are oriented 3’ to 5’ with the centromere located to the right (nucleotide coordinates are indicated). Percentages are calculated relative to total split reads mapping from the HO site to the last essential gene. Bars above the sequence correspond to the *UBP10* strain; bars below the sequence correspond to the *ubp10*Δ strain. In (A), a 33 bp region between Core 1 and Core 2 that is devoid of events is indicated with a gray triangle. (**C**) Percentages of total split reads mapping to SiRTA 9L-44 or a mutated SiRTA 9L-44 (Stim^mut^) are shown in *UBP10* and *ubp10*Δ strains. To generate 9L-44 Stim^mut^, the 52-bp Stim region of 9L-44 was replaced with a non-telomeric sequence. Data are from 3 pools of 30 GCR events. (**D**) Percentages of total split reads mapping to WT SiRTA 9L-44 or a mutated SiRTA 9L-44 (BS2^mut^) are shown in *UBP10* and *ubp10*Δ strains. To generate 9L-44 BS2^mut^, the 18-bp that compose SiRTA 9L-44 Cdc13 binding site 2 (BS2) were replaced with polyadenine. Data are from 2 pools of 30 GCR events. (**E**) Percentages of total split reads mapping to WT SiRTA 5L-35 or a mutated SiRTA 5L-35 (Stim^mut^) are shown in *UBP10* and *ubp10*Δ strains. To generate 5L-35 Stim^mut^, the 18-bp Stim region of SiRTA 5L-35 was replaced with polyadenine. Data are from 3 pools of 30 GCR events.

In addition to the previously identified SiRTA 9L-44, telomere addition is favored (>5% of total split reads) at two additional sites in the *UBP10* strain. The first site is 200 bp proximal to the HO cleavage site (41.6-41.7 kb), where more than 15% of GCR events occur (Figure 3A and 3B, arrows). In this region, the fraction of events involving telomere addition is strongly reduced by deletion of *UBP10*, with concomitant increase in translocations and deletions (Figure 3A and 3B; Supplement Figure 5A). The second site is between 42.2 and 42.4 kb from the endogenous telomere (Figure 3A and 3B, bracket). This site undergoes telomere addition in the *UBP10* strain, but incurs translocations and deletions in the absence of *UBP10* (Figure 3A and 3B; Supplement Figure 5B). To determine if these regions contain a TG-rich sequence similar to SiRTA 9L-44, we calculated the percentage of total nucleotides that are either T or G (T+G) and the ratio of G to T nucleotides in a sliding window along the strand running 3’ to 5’ between the HOcs and SiRTA 9L-44. In this region, SiRTA 9L-44 has the highest G/T ratio, but the 41.6-41.7 kb and 42.2-42.4 kb regions also have elevated G/T ratios compared to their surrounding sequences (Supplement Figure 5C). This analysis suggests that these sequences may function as SiRTAs.

Deletion of *UBP10* does not increase the frequency of translocations under all circumstances. The HOcs itself is a “hotspot” of *dn*TA, at least in part because cleavage generates a TGTT 3’ overhang upon which telomerase acts (Supplement Table 3). We compared the frequency of translocation split reads at the HOcs (41.5-41.6) and at SiRTA 9L-44 in the presence or absence of *UBP10.* In contrast to the striking increase in translocation events observed at SiRTA 9L-44, translocations do not significantly increase at the HOcs in the absence of *UBP10* (Figure 3C). The same result is observed on chromosome V (Supplement Figure 4C). Telomerase action at the HOcs is likely stimulated by the Ku heterodimer (Yku70/80), which associates with the DNA end to recruit telomerase through interaction with the telomerase RNA (36). These events appear unaffected by deletion of *UBP10*.

In summary, the patterns of GCR events incurred following a DSB on chromosome IX (Figure 3; Supplement Figure 5) and V (Supplement Figure 4) suggest that Ubp10 preferentially suppresses translocations at sites that are hotspots of *dn*TA, most prominently those at which *dn*TA is stimulated by the association of Cdc13 with single-stranded DNA following a DSB. These observations suggest that Cdc13 may stimulate chromosome rearrangements in the absence of *UBP10*, a possibility addressed in the next section.

### Association of Cdc13 with the SiRTA stimulates rearrangements in the absence of *UBP10*

We previously determined that SiRTAs 9L-44 and 5L-35 comprise a bipartite structure in which telomere addition at one or more Core sequences is promoted by a region (the “Stim”) located more distal to the chromosome break (24). We first examined how deletion of *UBP10* affects the precise location of *dn*TA and translocation events in the SiRTA. As previously reported, the majority of GCR events at SiRTA 9L-44 in the *UBP10* background involve telomere addition within the TG-rich Core 1 region, with a fraction of events occurring at Core 2, located ∼100 bp distal to Core 1 (Figure 4A) (24). With improved resolution, it is apparent that some telomere addition events occur between Core 1 and Core 2. In the absence of *UBP10*, *dn*TA events are reduced in frequency, but continue to cluster in Core 1. Translocation breakpoints predominantly occur in and near Core 1 (Figure 4A). At SiRTA 5L-35, *dn*TA events are clustered in the Core sequence in the *UBP10* strain, as previously reported (24). Upon deletion of *UBP10*, the overall pattern of *dn*TA is unchanged. In contrast to SiRTA 9L-44, translocations at SiRTA 5L-35 are not favored at the Core sequence, with most mapping proximal to the previously defined Stim sequence (Figure 4B).

Mutation of the Stim regions of SiRTAs 5L-35 or 9L-44 strongly reduces the frequency of *de novo* telomere addition by eliminating or reducing Cdc13 association (24). To test whether translocations observed in the absence of *UBP10* require Cdc13 binding, we replaced the entire Stim region of SiRTA 9L-44 (consisting of two Cdc13 binding sites, BS1 and BS2; Figure 4A) with a non-telomeric sequence (Stim^mut^). Importantly, this mutation does not alter sequences that engage in the majority of *dn*TA or translocation events (Figure 4A). As expected, Stim^mut^ significantly reduces *dn*TA in the *UBP10* strain (Figure 4C; Supplement Figure 6A). Strikingly, translocations previously observed in the *ubp10*Δ background are eliminated by Stim^mut^ (Figure 4C; Supplement Figure 6B). To verify this result, we mutated BS2 only, previously shown to eliminate *dn*TA at SiRTA 9L-44 (24). Indeed, mutation of BS2 eliminates translocations and *dn*TA events in the *ubp10*Δ strain (Figure 4D). It is unclear why the BS2 mutation has a stronger phenotype than Stim^mut^, but the difference may arise from sequences used to replace the Stim in each case. Mutating the Stim sequence at SiRTA 5L-35 in the *ubp10*Δ background eliminates translocations, although significance is difficult to ascertain since translocation frequencies are low at SiRTA 5L-35 (Figure 4E). Taken together, these results show that the Stim sequence is required to promote translocations at SiRTAs in the absence of *UBP10*, implicating Cdc13 in the stimulation of both *dn*TA and translocations.

If association of Cdc13 with the resected strand following a DSB is required to stimulate both *dn*TA and translocations (in the absence of *UBP10*), we reasoned that addition of a recognition site for Cdc13 to the extremely weak SiRTA 6L-22 might be sufficient to increase both types of events. *In vitro,* Cdc13 binds an 11-base single-stranded DNA oligonucleotide representative of yeast telomeric DNA sequence (Tel11; 5’-GTGTGGGTGTG-3’) with 3pM affinity (37). Integration of Tel11 upstream of the weakly functional SiRTA 6L-22 sequence dramatically increases the frequency of *de novo* telomere addition events at the SiRTA in the *UBP10* strain (from ∼6% to ∼85%; Figure 5A). Consistent with Tel11 serving as a Stim sequence, most *dn*TA events occur distal to Tel11 (Figure 5C). While the frequency of GCR events is unaffected by deletion of *UBP10*, repair shifts from *de novo* telomere addition to translocations, with translocations accounting for 44% of the total events at the mutated SiRTA (Figure 5A and 5B; Supplement Figure 7). Similar to the situation at SiRTA 5L-35, we observe translocations upstream of the Tel11 sequence, suggesting that Cdc13, bound to the single-stranded DNA produced by resection, can facilitate translocations even when the binding site is excluded from the product of the repair event (Figure 4B and Figure 5C). Taken together, these data strongly argue that Cdc13 is necessary and sufficient to stimulate translocations at SiRTAs in the absence of *UBP10*.

**Figure 5.**
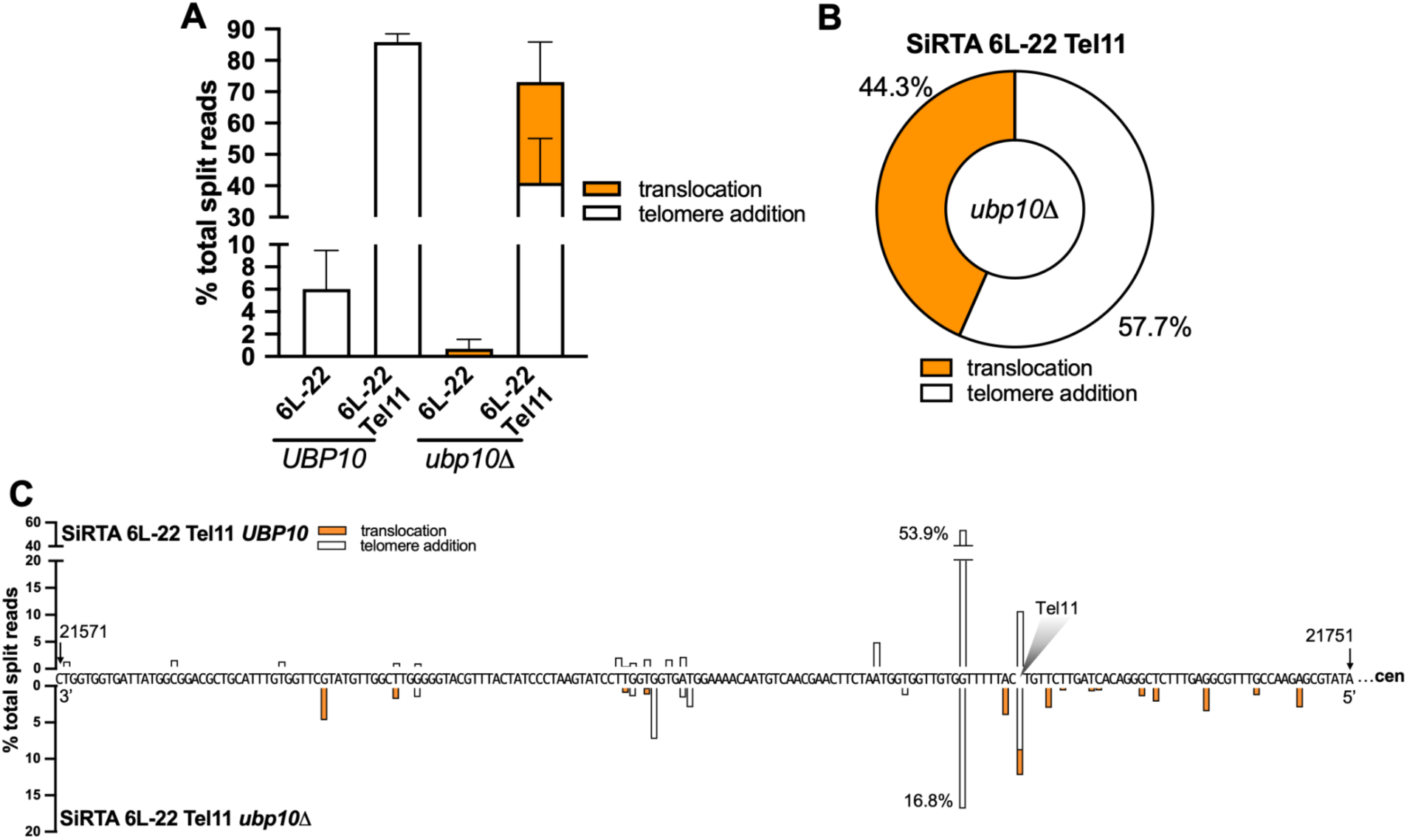
Integration of a single Cdc13 binding site is sufficient to stimulate translocations in the absence of *UBP10.* (**A-C**) DNA repair events were monitored across the region containing SiRTA 6L-22 and SiRTA 6L-22 Tel11 in *UBP10* and *ubp10*Δ strains. To generate SiRTA 6L-22 Tel11, the Tel11 sequence (5’-GTGTGGGTGTG-3’) was integrated upstream of SiRTA 6L-22. Data are from 3 pools of 30 GCR events each in each strain. (**A**) Percentages of total split reads that map to SiRTA 6L-22 or SiRTA 6L-22 Tel11 are shown. Data for SiRTA 6L-22 are repeated from Figure 2D for comparison. (**B**) Among split reads mapping to SiRTA 6L-22 Tel11, the percentages that involve telomere additions (white) or translocations (orange) are shown for the *ubp10*Δ strain. (**C**) Nucleotide-resolution maps showing sites of *de novo* telomere addition and/or translocations at SiRTA 6L-22 Tel11. The sequence is oriented 3’ to 5’ with the centromere located to the right. Bars above the sequence correspond to the *UBP10* strain; bars below the sequence correspond to the *ubp10*Δ strain. The shaded triangle represented the site at which the Tel11 sequence was integrated.

### Translocations at SiRTAs primarily target sub-telomeric sequences

Split reads contain not only the site within the SiRTA at which a translocation occurs, but also the sequence to which the SiRTA is joined (denoted here as the “target” sequence). The vast majority of translocations involve subtelomeric sequences that occur in *cis* or in *trans*. Subtelomeric translocations can be further subdivided into three classes (A-C) as described below.

Class A translocations, observed thus far only at SiRTA 9L-44, join the SiRTA to a site internal to either an X element or a Y’ element (Figure 6A; Supplement Figure 8 and 9A). Although X and Y’ elements show high levels of sequence identity, the split read sequences identified at SiRTA 9L-44 have only two identical matches in the reference genome (to X and Y’ elements on chromosome IX-L and X-L). The strain utilized in this work is from the S288C background and our prior analysis is consistent with high similarity to the reference strain (26). Therefore, Class A “translocations” are either large, internal deletions that join the SiRTA with the X or Y’ element on the left arm of chromosome IX (*cis*) or are true translocations that join the SiRTA with either the X or Y’ element on the left arm of chromosome X (*trans)*. Class A breakpoints occur at several closely clustered sequences at SiRTA 9L-44 (near Core 1; Figure 4A). Alignment of the template and donor sequences reveals, in each case, a region of interrupted microhomology (19-33 bp) at the translocation breakpoint (Supplement Figure 9A). The restriction of Class A events to SiRTA 9L-44 likely reflects the existence of fortuitous microhomology, perhaps combined with a higher likelihood of repair occurring in *cis* to the original DSB.

**Figure 6.**
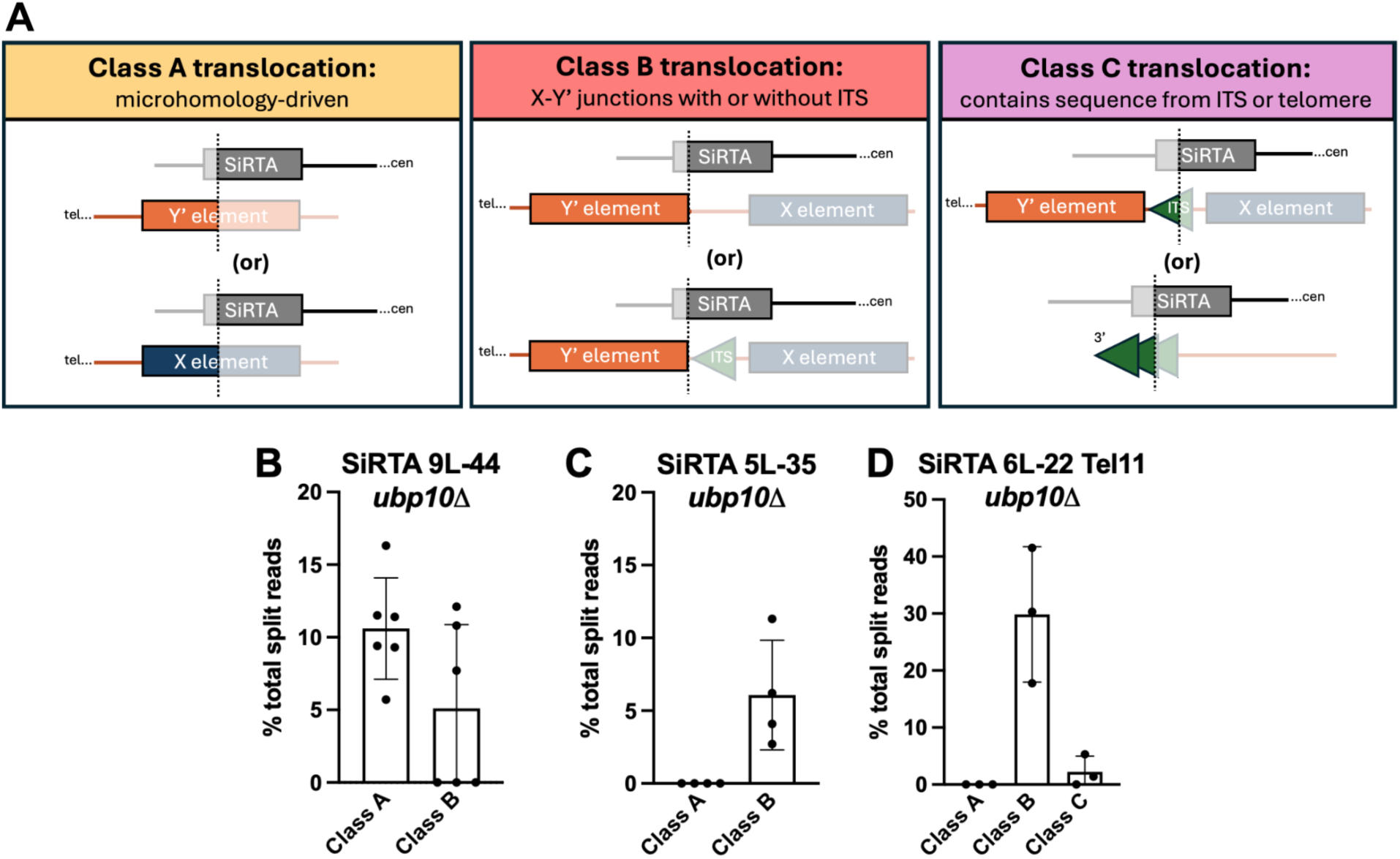
Ubp10 suppresses subtelomeric recombination at SiRTAs. (**A**) Schematic diagram depicting the classes of translocation breakpoints observed between the SiRTA (recipient; top) and the template chromosome (donor; bottom). Diagrams are oriented with the centromere to the right. Saturated colors illustrate the product observed in the translocation split read. The dotted line represents the translocation breakpoint. (**B-D**) Translocation split reads previously mapped to SiRTA 9L-44 (Figure 2A), SiRTA 5L-35 (Figure 2C), or SiRTA 6L-22 Tel11 (Figure 5A) were classified as Class A, Class B, or Class C and quantified as the percentage of total split reads between the HO cleavage site and the last essential gene. Each data point is generated from a pool of 30 GCR events. At SiRTA 5L-35, a fraction of translocations at the SiRTA is neither Class A nor B, with breakpoints in non-telomeric regions (average of 2% of total split reads).

Class B translocations join the SiRTA to a sequence found at the beginning of a subset of full-length Y’ elements (Figure 6A; Supplement Figure 8 and 9B). This is the most ubiquitous class of translocations, occurring at SiRTAs 9L-44, 5L-35, and the modified 6L-22 Tel11 (Figure 6B-D). The characteristic 5’-ATATATAT-3’ sequence at the breakpoint of the Class B translocations is found twelve times at subtelomeres, but the sequence located immediately proximal to this sequence motif varies (Supplement Figure 9B). In seven cases, the ATATATAT motif is immediately preceded by an ITS, while in the other five cases, the preceding sequence is unique or nearly unique. The subtelomeres of V-L and IX-L do not contain the ATATATAT motif, so the events observed at SiRTAs 5L-35 and 9L-44 must be translocations. The motif does occur on chromosome VI-L, so it is possible that at least some events at SiRTA 6L-22 Tel11 occur *in cis*.

Given the read length and high level of homology between X and Y’ elements, we cannot determine which X and Y’ elements are targeted (Supplement Figure 9B). However, there is no obvious sequence similarity between the SiRTA sequences that are involved in Class B translocations and little or no microhomology that could explain the exquisite targeting of these translocations to the first nucleotide of these particular Y’ elements (Supplement Figure 9B). For this reason, we favor the hypothesis that Class B translocations occur at one of the seven Y’ elements that are preceded by an ITS. Although SiRTAs contain telomere-like sequences, the 20 bp preceding the breakpoints of the Class B translocations are not remarkably TG-rich (Supplement Figure 9B), reinforcing the conclusion that Class B translocations are not microhomology mediated.

Class C translocations occur at SiRTA 14L-35 and are described above as events that resemble *dn*TA but require *RAD52* (Figure 3D and Figure 6A), suggesting a homology-mediated recombination event between SiRTA 14L-35 and either terminal or interstitial telomeric repeats. Rare Class C translocation split reads show evidence of Y’ sequence following the telomeric tract, consistent with at least a fraction of translocations occurring at an ITS (Supplement Figure 9C). Interestingly, although not perfectly homologous, the highest scoring match is the ITS on the left arm of chromosome XIV, consistent with a large deletion. Due to limitations of short-read sequencing, we are unable to quantify the relative use of the interstitial versus the terminal tract as a repair template. Our prior observation that deletion of *RAD52* fails to reduce telomere addition at SiRTAs 9L-44 or 5L-35 (26) suggests that Class C translocations are mostly limited to sites, like SiRTA 14L-35, that contain a long tract of sequence with high similarity to the telomeric TG_1-3_ pattern.

We observed two events at SiRTA 6L-22 Tel11 in which the ATATATAT motif is immediately preceded by TG-rich sequence that is not present at the SiRTA. In one case, the preceding sequence is 33 nucleotides of TG_1-3_ sequence, consistent with a Class C event (Supplement Figure 9C). Although imperfect, the closest match to this sequence is the ITS on the left arm of chromosome VI, suggesting again a preference for interactions in *cis* to the break. The other event contains “GTGT” and may represent a Class C event (there are two sites where this sequence precedes the ATATATAT motif). Alternatively, we cannot rule out the possibility that telomerase extended the 3’ end at the SiRTA as an intermediate step in repair.

At SiRTA 5L-35, ∼2% of total GCR events were translocation split reads that joined chromosome V to a non-telomeric sequence of a different chromosome arm. Some target sequences are located >100 kb from the nearest telomere and oriented in a way that would produce a dicentric chromosome. These breakpoints likely represent the initiation of a more complex rearrangement. It is unclear why this type of translocation is preferentially observed at SiRTA 5L-35. These non-subtelomeric translocations are not observed in the SiRTA 5L-35 Stim^mut^ *ubp10*Δ strain, suggesting that they depend on Cdc13 association. Cdc13 binding is not sufficient, however, since no events of this type are observed at SiRTA 6L-22 Tel11.

### Ubp10 represses homologous recombination repair mechanisms at SiRTAs

We examined the requirements for translocations/deletions at SiRTA 9L-44 in the context of *UBP10* loss. As expected, deletion of *RAD52*, broadly required for homologous recombination, eliminates chromosome rearrangements at the SiRTA (Figure 7A). In contrast, chromosome rearrangements at SiRTA 9L-44 are reduced, but not eliminated, by deletion of *RAD51* (Figure 7A). Upon closer inspection, the remaining rearrangements in the *ubp10*Δ *rad51*Δ strain are exclusively Class B, suggesting that Class A events require Rad51, but Class B do not (Figure 7B). We confirmed this result at SiRTA 6L-22 Tel11, where nearly all rearrangements in the absence of *UBP10* are Class B, accounting for ∼30% of total GCR events (Figure 6D). While the frequency is reduced upon additional deletion of *RAD51* (to about 40% of that in the *RAD51 ubp10*Δ strain), Class B translocations clearly persist (Figure 7C and Supplement Figure 10B), confirming that these events can occur through a *RAD51*-independent mechanism. Both Class A and Class B events at SiRTA 9L-44 require *RAD59*, but are unaffected by deletion of *RAD54* (Figure 7A and B). As expected, *dn*TA events persist at SiRTA 9L-44 in the absence of homologous recombination (Supplement Figure 10A), although *dn*TA events are rare after deletion of *RAD51*, as we previously reported (26).

**Figure 7.**
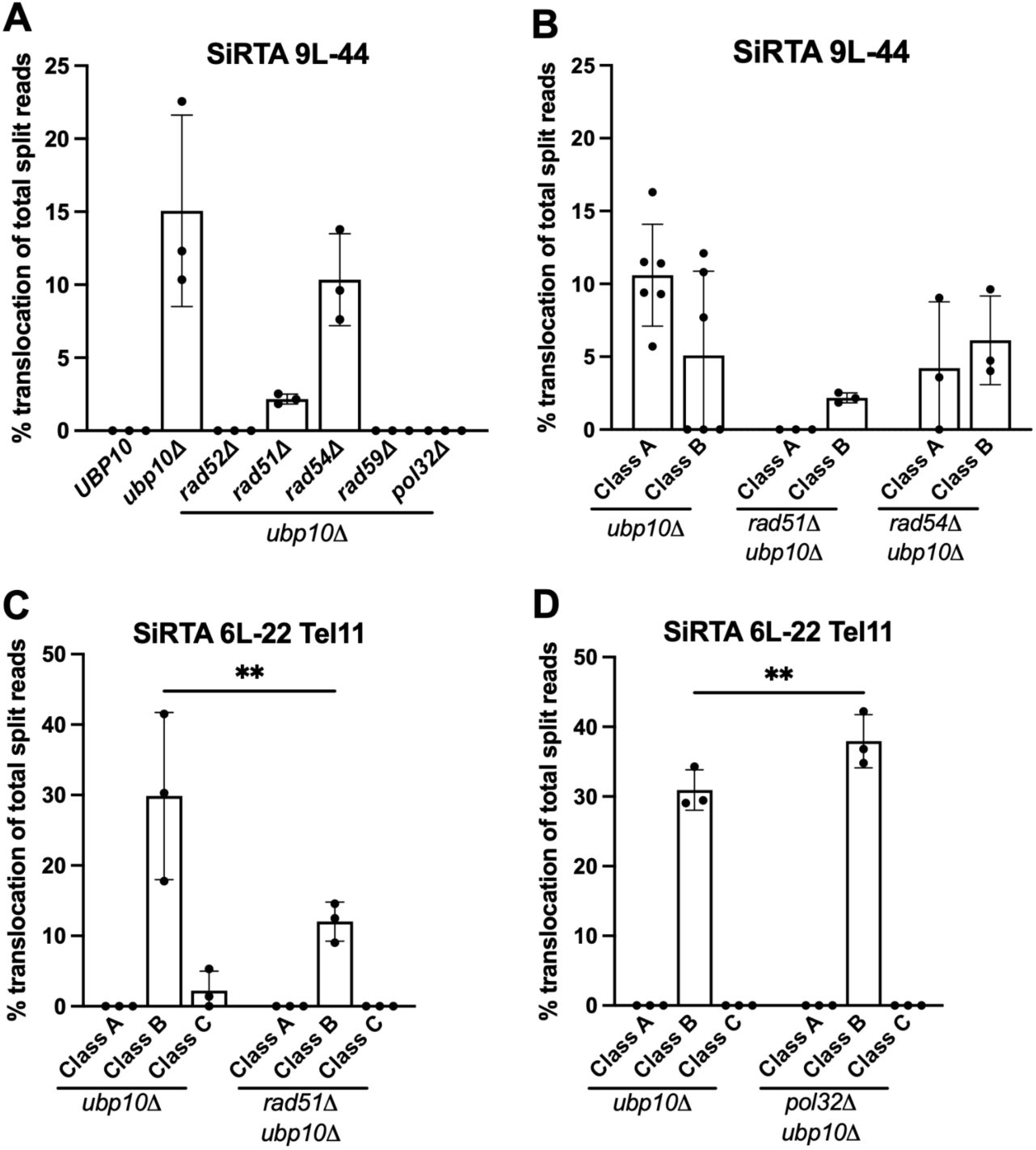
Class B translocations occur independently of *RAD51* and *POL32*. (**A**) The percentage of translocation split reads out of total split reads is shown at SiRTA 9L-44 for each strain indicated. Data are from 3 pools of 30 GCR events. (**B**) Of the strains that exhibited translocation events at SiRTA 9L-44, translocation split reads were assigned as Class A or Class B translocations. Class C translocations were not observed. Data for the *ubp10*Δ strain are repeated from Figure 6B for comparison. (**C**) Translocation split reads were assigned to classes at SiRTA 6L-22 Tel11 in the *rad51*Δ *ubp10*Δ mutant. Data for the *ubp10*Δ strain are repeated from Figure 6D for comparison. \*\**p*<0.005, Two-way ANOVA with Šídák’s multiple comparisons test. (**D**) Translocation split reads were divided in classes at SiRTA 6L-22 Tel11 in the *pol32*Δ *ubp10*Δ mutant. \*\**p*<0.005 Two-Way ANOVA with Šídák’s multiple comparisons test.

In the experiments described here, we select for clones that have lost sequences distal to the HO cleavage site. Therefore, true translocations recovered in this assay are non-reciprocal and likely arise through break-induced replication (BIR), a mechanism that operates when only one end of a DSB can productively engage in repair (13, 38, 39). The requirement for *RAD52* is consistent with, but not diagnostic of, this type of repair. To evaluate the involvement of BIR, we deleted the nonessential subunit of Polδ (encoded by *POL32*), which is required for successful completion of BIR (40, 41). At SiRTA 9L-44, Class A and Class B events are abolished in the *pol32*Δ *ubp10*Δ strain (Figure 7A). For Class B events, this result is expected since the donor sequence is not present on the left telomere of chromosome IX (i.e. these must be translocations). In contrast, Class A events likely involve repair between SiRTA 9L-44 and the chromosome IX subtelomere, although we cannot rule out translocations with identical sequences on chromosome X. The requirement for Pol32 indicates that Class A events occur through BIR rather than single-strand annealing (SSA), with invasion and duplication of distal sequences occurring in *cis* prior to degradation of the chromosome terminus (13, 40, 42). A similar preference for BIR between repeated sequences has been observed in mammalian cells when the break is located asymmetrically between the repeats (13, 39).

It is surprising that Class A events require Rad51 since events associated with limited microhomology have been previously associated with Rad51-independent BIR (13, 43–45). We speculate that the requirement for Rad51 reflects its role in promoting Cdc13 association with SiRTAs, rather than a direct effect on recombination. We previously found that deletion of *RAD51* impairs *dn*TA at SiRTAs and is associated with reduced recruitment of Cdc13 to SiRTA 5L-35 following a DSB (26). Given that Cdc13 stimulates rearrangements at SiRTAs (Figures 4, 5), the decreased association of Cdc13 that accompanies loss of *RAD51* is predicted to reduce events in the *ubp10*Δ background. Class B events persist in the absence of *RAD51*, suggesting that they may be more tolerant of reduced Cdc13 binding.

Given that Pol32 appears to be required for all events at SiRTA 9L-44, we were surprised to find that Class B events at SiRTA 6L-22 Tel11 persist in the *pol32*Δ *ubp10*Δ background (Figure 7D). It is unclear whether there is something different about the Class B events on chromosome VI compared to those on chromosome IX or whether the high efficiency of events on chromosome VI masks a partial requirement for Pol32. Pol32 is not required for the initiation of BIR and synthesis can continue as much as 15 kb in its absence (41). Given the subtelomeric locations of the target sequence for Class B events, replication extends 4.5 to 7 kb to reach the chromosome end and therefore may not rely strongly on Pol32. The chromosome VI left subtelomere contains an ITS followed by the 5’-ATATATAT-3’ motif utilized in Class B events. Therefore, it is also possible that at least a subset of events on chromosome VI occur through SSA, which would be expected to require *RAD52* but be independent of *RAD51* (15).

### Sir4 and Sir2 are required for a subset of subtelomeric translocations

The observation that most rearrangements in the *ubp10*Δ strain involve subtelomeric sequences suggests a potential mechanism for Ubp10 action. Yeast subtelomeric regions are transcriptionally silenced by recruitment of the Silent Information Regulator (SIR) proteins (46, 47). Ubp10 physically interacts with Sir4 to deubiquitinate H2B, promoting heterochromatin formation at telomeric loci (48, 49). In the absence of *UBP10*, disruption of subtelomeric heterochromatin may provide a favorable substrate for recombination. If true, deletion of *SIR4* should promote rearrangements at SiRTAs, similar to deletion of *UBP10*. In contrast, we observe only *dn*TA events at SiRTA 9L-44 in the absence of *SIR4* (Supplement Figure 10C), indicating that loss of telomeric silencing alone does not explain the phenotype of the *ubp10*Δ strain.

As a control, we deleted *SIR4* in the *ubp10*Δ background. Although the overall frequency of rearrangements at SiRTA 9L-44 was unchanged, Class B events were specifically eliminated (Figure 8A, B). To verify this result, we examined the effect of deleting *UBP10* and *SIR4* at SiRTA 6L-22 Tel11, where Class B translocations are prominent. As expected, rearrangements at SiRTA 6L-22 are eliminated in the absence of *SIR4* (Figure 8C), although *dn*TA is unaffected (Supplement Figure 10D). Sir4 promotes telomeric silencing as part of the SIR complex that also includes Sir2 and Sir3. We observed a loss of rearrangements in the absence of *SIR2* at SiRTA 6L-22 Tel11; however, the frequency and type of events remained unaffected in the absence of *SIR3* (Figure 8C, D; Supplement Figure 10D), indicating that the role of the SIR complex in telomeric silencing cannot explain the effect on Class B translocations. Loss of silencing at the mating type loci can indirectly affect DNA repair pathways by activating gene expression patterns typical of diploid cells (50). However, in addition to the lack of effect with *SIR3* deletion, the strains that we use for these experiments contain deletions at the silent mating loci that further rule out this potential explanation for the loss of Class B translocations upon deletion of *SIR2* or *SIR4*.

**Figure 8.**
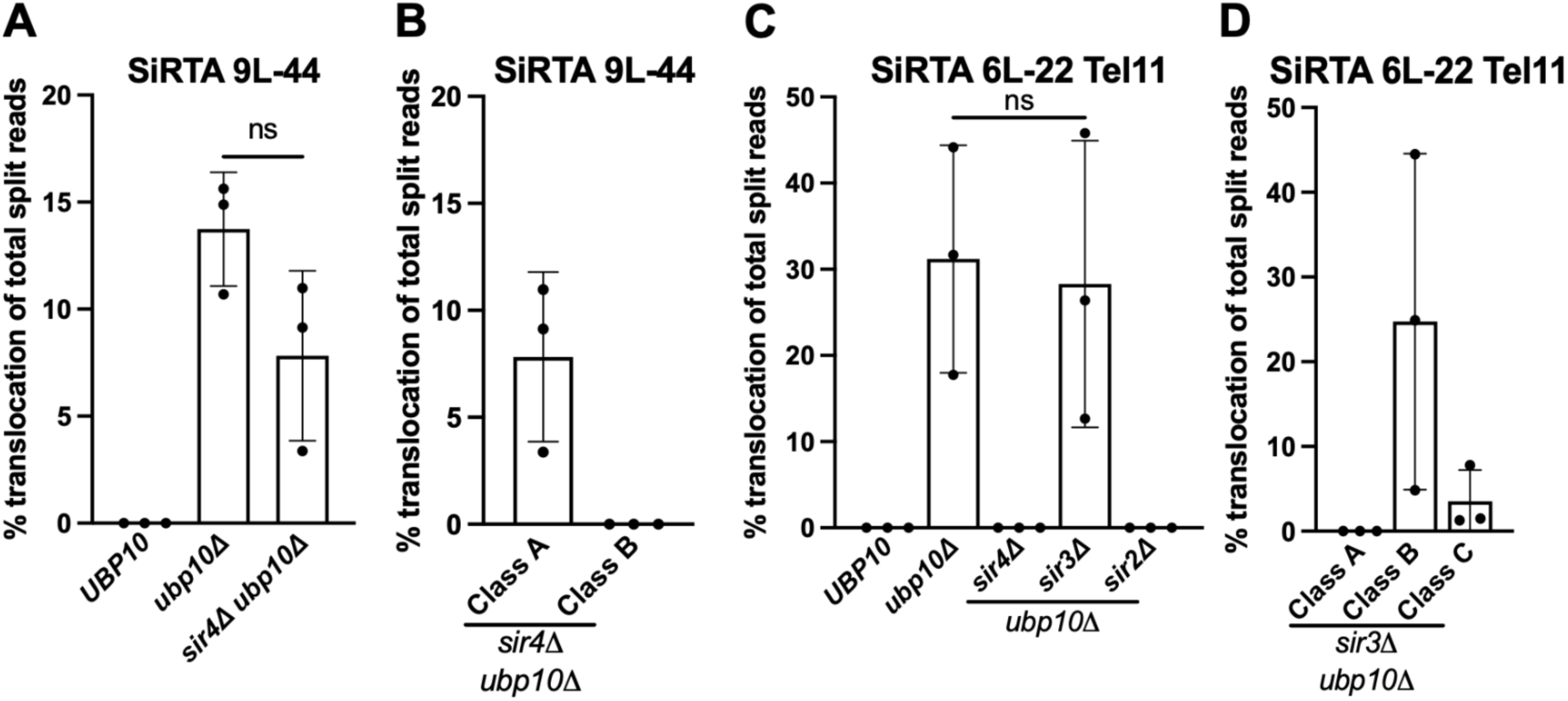
Class B translocations require Sir4 and Sir2. (**A**) Percentages of translocation split reads (of total split reads) are shown for the indicated strains. Data are from 3 pools of 30 GCR events. \**p*<0.05 by ANOVA with Dunnett’s multiple comparisons test. (**B**) Translocation split reads observed at SiRTA 9L-44 in the *sir4*Δ *ubp10*Δ strain were divided into their respective classes. (**C**) Percentages of translocation split reads (of total split reads) are shown at SiRTA 6L-22 Tel11 for the indicated strains. Data are from 3 pools of 30 GCR events. Data for the *ubp10*Δ strain are from Figure 7C. (**D**) Translocation split reads observed at SiRTA 6L-22 Tel11 in the *sir3*Δ *ubp10*Δ strain were divided into their respective classes.

Sir4 plays a key role in anchoring subtelomeric chromosomal regions to the nuclear periphery through interactions with nuclear membrane proteins (51–53). Given that persistent DNA breaks are recruited near the nuclear envelope, it is possible that translocations are promoted by the spatial proximity between the Cdc13-bound SiRTA sequence and the subtelomeric ITS within the same nuclear subcompartment (52, 54). The differential effects of Sir2 and Sir3 are less easily explained, although Sir3 has been reported to be dispensable for the localization of telomeres to the nuclear periphery (55). It is possible that Sir2 (in complex with Sir4) may help stabilize the association of the broken end with the nuclear periphery and with subtelomeric chromatin, creating an environment conducive for recombination with telomeric sequences. We note that the differential dependencies of Class A and Class B translocations on Rad51, Sir4, and Sir2 suggest that these rearrangements arise through different mechanisms, despite both using subtelomeric sequence as templates.

### Conclusions and implications

The observations described here show that association of Cdc13 with telomere-like sequences following a DSB promotes subtelomeric rearrangements in a manner that is normally suppressed by the deubiquitinase activity of Ubp10 (Figure 2B). Ubp10 targets ubiquitinated histone 2B (H2B) and proliferating cell nuclear antigen (PCNA), both of which are associated with DNA damage signaling pathways (56–61). While loss of telomeric silencing alone does not explain the phenotype of the *ubp10*Δ strain (Figure 7F), we have not ruled out the possibility that rearrangements are facilitated by increased levels of ubiquitinated H2B. Mono- or poly-ubiquitinated PCNA promotes translesion synthesis (TLS) and template switching (TS), respectively (58, 62). Furthermore, Ubp10 action promotes unloading of PCNA during lagging strand replication, required for normal Okazaki fragment processing (63). Failure to appropriately regulate PCNA ubiquitination may promote rearrangements by facilitating either the initiation or the successful completion of BIR within subtelomeric sequences. For example, Rad51-independent BIR is predicted to require generation of single-stranded DNA to circumvent the need for the strand invasion activity of Rad51 (16, 39, 64). Alternatively, promotion of template switching activities may facilitate successful completion of BIR-mediated repair.

The translocations observed at SiRTAs resemble survivors that utilize the alternative lengthening of telomeres (ALT) recombination mechanism used in 10-15% of cancer cells to maintain their telomeres (17, 18). Originally characterized in *S. cerevisiae,* budding yeast has served as a robust model to study ALT. However, analysis of initial ALT survivor formation remains challenging due to the genomic instability generated in cells lacking telomerase. Here, we identified a unique situation that allows us to probe the mechanism of ALT-like survivor formation specifically with the events observed at SiRTA 14L-35 (Figure 2E).

In this work, we demonstrated that association of Cdc13 following a DSB can stimulate the formation of translocations, suggesting a previously unanticipated role of Cdc13 in facilitating events involving telomeric and telomere-like sequences (Figures 4, 5). Tethering of persistent DSBs to the nuclear periphery relies on interactions between subtelomeric binding proteins and nuclear envelope components (52, 65, 66). Cdc13 has been suggested to promote the sequestration of persistent DSBs through its association with the telomerase component Est1 (52). Most translocations observed at SiRTAs contain no telomeric sequence at the translocation junction, suggesting that synthesis by telomerase is not required to stimulate these events. We speculate that Cdc13 may contribute to repair of DSBs at SiRTAs by promoting the colocalization of breaks and telomerase/subtelomeric sequences. We observe events in which BIR is initiated both *in cis* and *in trans* to the HO-generated break, suggesting that the propensity of telomeres to cluster may contribute to these events. Together, these observations highlight the intricate interplay between telomere maintenance mechanisms and DNA damage repair pathways.

## DATA AVAILABILITY

Strains and plasmids are available upon request. The data presented in this article are available in the article, in its online Supplementary material, or will be shared upon reasonable request to the corresponding author. Sequencing data are available from the NIH Sequence Read Archive (SRA) under BioProject ID: PRJNA1279155.

## SUPPLEMENTARY DATA

Supplementary Data are available.

## ACKNOWLEDGEMENTS

We thank James Haber for providing us with strains and plasmids. We are grateful to the Vanderbilt VANTAGE sequencing core for performing the Illumina sequencing in this work.

## AUTHOR CONTRIBUTIONS

David Gonzalez: Conceptualization, Formal Analysis, Methodology, Validation, Visualization, Writing-original draft.

Allison Westerbeek: Methodology.

Esther Epum: Conceptualization, Formal Analysis, Methodology.

Katherine Friedman: Conceptualization, Formal Analysis, Methodology, Validation, Visualization, Writing-review & editing.

## REFERENCES

1. Blackburn, E.H. (1991) Structure and function of telomeres. Nature, 350, 569–573.

2. Wellinger, R.J. and Zakian, V.A. (2012) Everything you ever wanted to know about *Saccharomyces cerevisiae* telomeres: Beginning to end. Genetics, 191, 1073–1105.

3. Osterhage, J.L. and Friedman, K.L. (2009) Chromosome end maintenance by telomerase. Journal of Biological Chemistry, 284, 16061–16065.

4. Greider, C.W. and Blackburnt, E.H. A telomeric sequence in the RNA of *Tetrahymena* telomerase required for telomere repeat synthesis. Nature, 337, 331–337.

5. Pfeiffer, V. and Lingner, J. (2013) Replication of telomeres and the regulation of telomerase. Cold Spring Harb Perspect Biol, 5, a010405.

6. Pennock, E., Buckley, K. and Lundblad, V. (2001) Cdc13 delivers separate complexes to the telomere for end protection and replication. Cell, 104, 387–396.

7. Evans, S.K. and Lundblad, V. (1999) Est1 and Cdc13 as comediators of telomerase access. Science (1979), 286, 117–120.

8. Mehta, A. and Haber, J.E. (2014) Sources of DNA double-strand breaks and models of recombinational DNA repair. Cold Spring Harb Perspect Biol, 6, a016428.

9. Wang, X. and Haber, J.E. (2004) Role of *Saccharomyces* single-stranded DNA-binding protein RPA in the strand invasion step of double-strand break repair. PLoS Biol, 2, 0104–0112.

10. Ruff, P., Donnianni, R.A., Glancy, E., Oh, J. and Symington, L.S. (2016) RPA stabilization of single-stranded DNA is critical for break-induced replication. Cell Rep, 17, 3359–3368.

11. New, J.H., Sugiyama, T., Zaitseva, E. and Kowalczykowski, S.C. (1998) Rad52 protein stimulates DNA strand exchange by Rad51 and replication protein A. Nature, 391, 407–410.

12. Sung, P., Krejci, L., Van Komen, S. and Sehorn, M.G. (2003) Rad51 recombinase and recombination mediators. Journal of Biological Chemistry, 278, 42729–42732.

13. Malkova, A. (2018) Break-induced replication: The where, The why, and The how. Trends in Genetics, 34, 518–531.

14. McVey, M. and Lee, S.E. (2008) MMEJ repair of double-strand breaks (director’s cut): deleted sequences and alternative endings. Trends in Genetics, 24, 529–538.

15. Ivanov, E.L., Sugawara, N., Fishman-Lobell, J. and Haber, J.E. (1996) Genetic requirements for the single-strand annealing pathway of double-strand break repair in *Saccharomyces cerevisiae*. Genetics, 142, 693–704.

16. Malkova, A., Signon, L., Schaefer, C.B., Naylor, M.L., Theis, J.F., Newlon, C.S. and Haber, J.E. (2001) Rad51-independent break-induced replication to repair a broken chromosome depends on a distant enhancer site. Genes Dev, 15, 1055–1060.

17. Lundblad, V. and Blackburn, E.H. (1993) An alternative pathway for yeast telomere maintenance rescues *est1^−^* senescence. Cell, 73, 347–360.

18. Kockler, Z.W., Comeron, J.M. and Malkova, A. (2021) A unified alternative telomere-lengthening pathway in yeast survivor cells. Mol Cell, 81, 1816–1829.e5.

19. Kim, S., Park, S.H., Kang, N., Ra, J.S., Myung, K. and Lee, K. (2024) Polyubiquitinated PCNA triggers SLX4-mediated break-induced replication in alternative lengthening of telomeres (ALT) cancer cells. Nucleic Acids Res, 52, 11785–11805.

20. Putnam, C.D., Pennaneach, V. and Kolodner, R.D. (2004) Chromosome healing through terminal deletions generated by *de novo* telomere additions in *Saccharomyces cerevisiae*. PNAS, 101, 13262–13267.

21. Pennaneach, V., Putnam, C.D. and Kolodner, R.D. (2006) Chromosome healing by *de novo* telomere addition in *Saccharomyces cerevisiae*. Mol Microbiol, 59, 1357–1368.

22. Stellwagen, A.E., Haimberger, Z.W., Veatch, J.R. and Gottschling, D.E. (2003) Ku interacts with telomerase RNA to promote telomere addition at native and broken chromosome ends. Genes Dev, 17, 2384–2395.

23. Myung, K., Datta, A. and Kolodner, R.D. (2001) Suppression of spontaneous chromosomal rearrangements by S phase checkpoint functions in *Saccharomyces cerevisiae*. Cell, 104, 397–408.

24. Obodo, U.C., Epum, E.A., Platts, M.H., Seloff, J., Dahlson, N.A., Velkovsky, S.M., Paul, S.R. and Friedman, K.L. (2016) Endogenous hot spots of *de novo* telomere addition in the yeast genome contain proximal enhancers that bind Cdc13. Mol Cell Biol, 36, 1750–1763.

25. Ngo, K., Gittens, T.H., Gonzalez, D.I., Hatmaker, E.A., Plotkin, S., Engle, M., Friedman, G.A., Goldin, M., Hoerr, R.E., Eichman, B.F., et al. (2023) A comprehensive map of hotspots of *de novo* telomere addition in *Saccharomyces cerevisiae*. Genetics, 224.

26. Epum, E.A., Mohan, M.J., Ruppe, N.P. and Friedman, K.L. (2020) Interaction of yeast Rad51 and Rad52 relieves Rad52-mediated inhibition of *de novo* telomere addition. PLoS Genet, 16, e1008608.

27. Lydeard, J.R., Lipkin-Moore, Z., Jain, S., Eapen, V. V. and Haber, J.E. (2010) Sgs1 and Exo1 redundantly inhibit break-induced replication and *de novo* telomere addition at broken chromosome ends. PLoS Genet, 6, e1000973.

28. Longtine, M.S., McKenzie, A., Demarini, D.J., Shah, N.G., Wach, A., Brachat, A., Philippsen, P. and Pringle, J.R. (1998) Additional modules for versatile and economical PCR-based gene deletion and modification in *Saccharomyces cerevisiae*. Yeast, 14, 953–961.

29. Sikorski, R.S. and Hieter, P. (1989) A system of shuttle vectors and yeast host strains designed for efficient manipulation of DNA in Saccharomyces cerevisiae. Genetics, 122, 19–27

30. Ngo, K., Epum, E.A. and Friedman, K.L. (2020) Emerging non-canonical roles for the Rad51– Rad52 interaction in response to double-strand breaks in yeast. Curr Genet, 66, 917–926.

31. Bolger, A.M., Lohse, M. and Usadel, B. (2014) Trimmomatic: A flexible trimmer for Illumina sequence data. Bioinformatics, 30, 2114–2120.

32. Langmead, B., Trapnell, C., Pop, M. and Salzberg, S.L. (2009) Ultrafast and memory-efficient alignment of short DNA sequences to the human genome. Genome Biol, 10, R25.

33. Langmead, B. and Salzberg, S.L. (2012) Fast gapped-read alignment with Bowtie 2. Nat Methods, 9, 357–359.

34. Robinson, D.E., Vanacloig-Pedros, E., Cai, R., Place, M., Hose, J. and Gasch, A.P. (2023) Gene-by-environment interactions influence the fitness cost of gene copy-number variation in yeast. G3: Genes, Genomes, Genetics, 13, jkad159.

35. Hoerr, R.E., Ngo, K. and Friedman, K.L. (2021) When the ends justify the means: Regulation of telomere addition at double-strand breaks in yeast. Front Cell Dev Biol, 9, 1–7.

36. Chen, H., Xue, J., Churikov, D., Hass, E.P., Shi, S., Lemon, L.D., Luciano, P., Bertuch, A.A., Zappulla, D.C., Géli, V., et al. (2018) Structural insights into yeast telomerase recruitment to telomeres. Cell, 172, 331–343.e13.

37. Eldridge, A.M. and Wuttke, D.S. (2008) Probing the mechanism of recognition of ssDNA by the Cdc13-DBD. Nucleic Acids Res, 36, 1624–1633.

38. Llorente, B., Smith, C.E. and Symington, L.S. (2008) Break-induced replication: What is it and what is it for? Cell Cycle, 7, 859–864.

39. Lee, R.S., Twarowski, J.M. and Malkova, A. (2024) Stressed? Break-induced replication comes to the rescue! DNA Repair (Amst), 142, 103759.

40. Lydeard, J.R., Jain, S., Yamaguchi, M. and Haber, J.E. (2007) Break-induced replication and telomerase-independent telomere maintenance require Pol32. Nature, 448, 820–823.

41. Liu, L., Yan, Z., Osia, B.A., Twarowski, J., Sun, L., Kramara, J., Lee, R.S., Kumar, S., Elango, R., Li, H., et al. (2021) Tracking break-induced replication shows that it stalls at roadblocks. Nature, 590, 655–659.

42. Deem, A., Keszthelyi, A., Blackgrove, T., Vayl, A., Coffey, B., Mathur, R., Chabes, A. and Malkova, A. (2011) Break-induced replication is highly inaccurate. PLoS Biol, 9, 1000594.

43. Sugawara, N., Ira, G. and Haber, J.E. (2000) DNA length dependence of the Single-Strand Annealing pathway and the role of *Saccharomyces cerevisiae* Rad59 in double-strand break repair. Mol Cell Biol, 20, 5300–5309.

44. Kockler, Z.W., Osia, B., Lee, R., Musmaker, K. and Malkova, A. (2021) Repair of DNA breaks by break-induced replication. Annu Rev Biochem, 90, 165–191.

45. Davis, A.P. and Symington, L.S. (2004) Rad51-dependent break-induced replication in yeast. Mol Cell Biol, 24, 2344–2351.

46. Aparicio, O.M., Billington, B.L. and Gottschling, D.E. (1991) Modifiers of position effect are shared between telomeric and silent mating-type loci in *S. cerevisiae*. Cell, 66, 1279–1287.

47. Hecht, A., Laroche, T., Strahi-Bolsinger, S., Gasser, S.M. and Grunstein, M. (1995) Histone H3 and H4 N-termini interact with Sir3and Sir4 proteins: A molecular model for the formation of heterochromatin in yeast.

48. Zukowski, A., Al-Afaleq, N.O., Duncan, E.D., Yao, T. and Johnson, A.M. (2018) Recruitment and allosteric stimulation of a histone-deubiquitinating enzyme during heterochromatin assembly. Journal of Biological Chemistry, 293, 2498–2509.

49. Kahana, A. and Gottschling, D.E. (1999) Dot4 links silencing and cell growth in *Saccharomyces cerevisiae*. Mol Cell Biol, 19, 6608–6620.

50. Kaeberlein, M., Mcvey, M. and Guarente, L. (1999) The Sir2/3/4 complex and Sir2 alone promote longevity in *Saccharomyces cerevisiae* by two different mechanisms. Genes Dev.

51. Lapetina, D.L., Ptak, C., Roesner, U.K. and Wozniak, R.W. (2017) Yeast silencing factor Sir4 and a subset of nucleoporins form a complex distinct from nuclear pore complexes. Journal of Cell Biology, 216, 3145–3159.

52. Oza, P., Jaspersen, S.L., Miele, A., Dekker, J. and Peterson, C.L. (2009) Mechanisms that regulate localization of a DNA double-strand break to the nuclear periphery. Genes Dev, 23, 912–927.

53. Therizols, P., Fairhead, C., Cabal, G.G., Genovesio, A., Olivo-Marin, J.C., Dujon, B. and Fabre, E. (2006) Telomere tethering at the nuclear periphery is essential for efficient DNA double strand break repair in subtelomeric region. Journal of Cell Biology, 172, 189–199.

54. Kalocsay, M., Hiller, N.J. and Jentsch, S. (2009) Chromosome-wide Rad51 spreading and Sumo-H2A.Z-dependent chromosome fixation in response to a persistent DNA double-strand break. Mol Cell, 33, 335–343.

55. Tham, W.-H., Stuart, J., Wyithe, B., Ferrigno, P.K., Silver, P.A. and Zakian, V.A. (2001) Localization of yeast telomeres to the nuclear periphery is separable from transcriptional repression and telomere stability functions. Mol Cell, 8, 189–199

56. Korenfeld, H.T., Avram-Shperling, A., Zukerman, Y., Iluz, A., Boocholez, H., Ben-Shimon, L. and Ben-Aroya, S. (2022) Reversal of histone H2B mono-ubiquitination is required for replication stress recovery. DNA Repair (Amst), 119, 103387.

57. Gardner, R.G., Nelson, Z.W. and Gottschling, D.E. (2005) Ubp10/Dot4p regulates the persistence of ubiquitinated histone H2B: Distinct roles in telomeric silencing and general chromatin. Mol Cell Biol, 25, 6123–6139.

58. Gallego-Sánchez, A., Andrés, S., Conde, F., San-Segundo, P.A. and Bueno, A. (2012) Reversal of PCNA ubiquitylation by Ubp10 in *Saccharomyces cerevisiae*. PLoS Genet, 8, e1002826.

59. Alvarez, V., Frattini, C., Sacristán, M.P., Gallego-Sánchez, A., Bermejo, R. and Bueno, A. (2019) PCNA deubiquitylases control DNA damage bypass at replication forks. Cell Rep, 29, 1323–1335.

60. Alvarez, V., Vinas, L., Gallego-Sanchez, A., Andres, S., Sacristan, M.P. and Bueno, A. (2016) Orderly progression through S-phase requires dynamic ubiquitylation and deubiquitylation of PCNA. Sci Rep, 6, 1–14.

61. Zamarreño, J., Muñoz, S., Alonso-Rodríguez, E., Alcalá, M., Rodríguez, S., Bermejo, R., Sacristán, M.P. and Bueno, A. (2024) Timely lagging strand maturation relies on Ubp10 deubiquitylase-mediated PCNA dissociation from replicating chromatin. Nat Commun, 15, 8183.

62. Masuda, Y. and Masutani, C. (2019) Spatiotemporal regulation of PCNA ubiquitination in damage tolerance pathways. Crit Rev Biochem Mol Biol, 54, 418–442.

63. Da Nguyen, H., Becker, J., Thu, Y.M., Costanzo, M., Koch, E.N., Smith, S., Myung, K., Myers, C.L., Boone, C. and Bielinsky, A.K. (2013) Unligated okazaki fragments induce PCNA ubiquitination and a requirement for Rad59-Dependent replication fork progression. PLoS One, 8.

64. Zhang, T., Rawal, Y., Jiang, H., Kwon, Y., Sung, P. and Greenberg, R.A. (2023) Break-induced replication orchestrates resection-dependent template switching. Nature, 619, 201–208.

65. Strambio-De-Castillia, C., Niepel, M. and Rout, M.P. (2010) The nuclear pore complex: Bridging nuclear transport and gene regulation. Nat Rev Mol Cell Biol, 11, 490–501.

66. Mekhail, K. and Moazed, D. (2010) The nuclear envelope in genome organization, expression and stability. Nat Rev Mol Cell Biol, 11, 317–328.

