## Supplement Figures for "The ubiquitin protease Ubp10 suppresses the formation of translocations at Cdc13 binding sites"

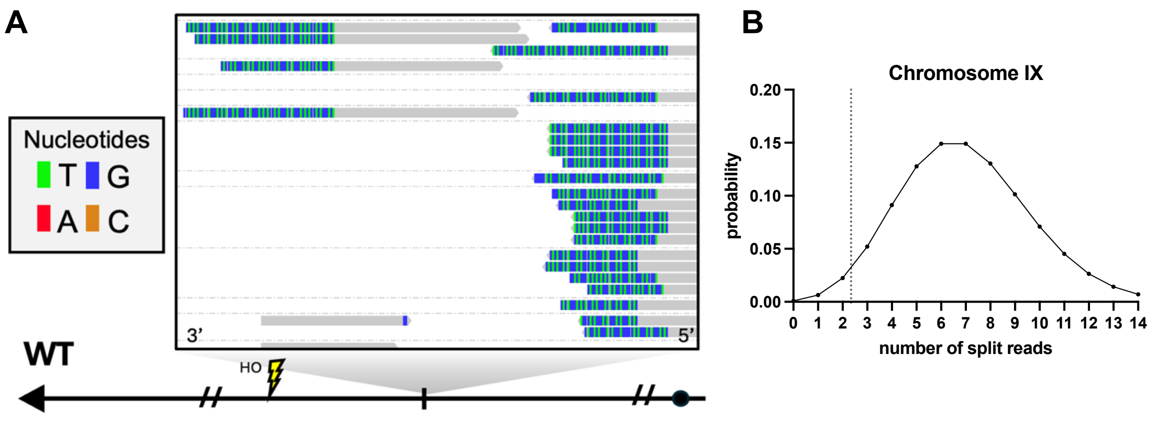


**Supplement Figure 1.** (**A**) Representative image of telomere addition split reads that contain $TG_{1\text{-}3}$ or $CA_{1\text{-}3}$ nucleotides at SiRTA 9L-44. (**B**) Poisson distribution model predicting the expected number of split reads per gross chromosomal rearrangement (GCR) event. A normal distribution was assumed given an average of 127 split reads in a sample of 30 independent GCR events. The number of split reads expected from a single GCR event was predicted based on this model. 95% of events were predicted to be represented by 2.4 or more GCR split reads (vertical dotted line). Those represented by at least two split reads were considered a bonafide GCR event.

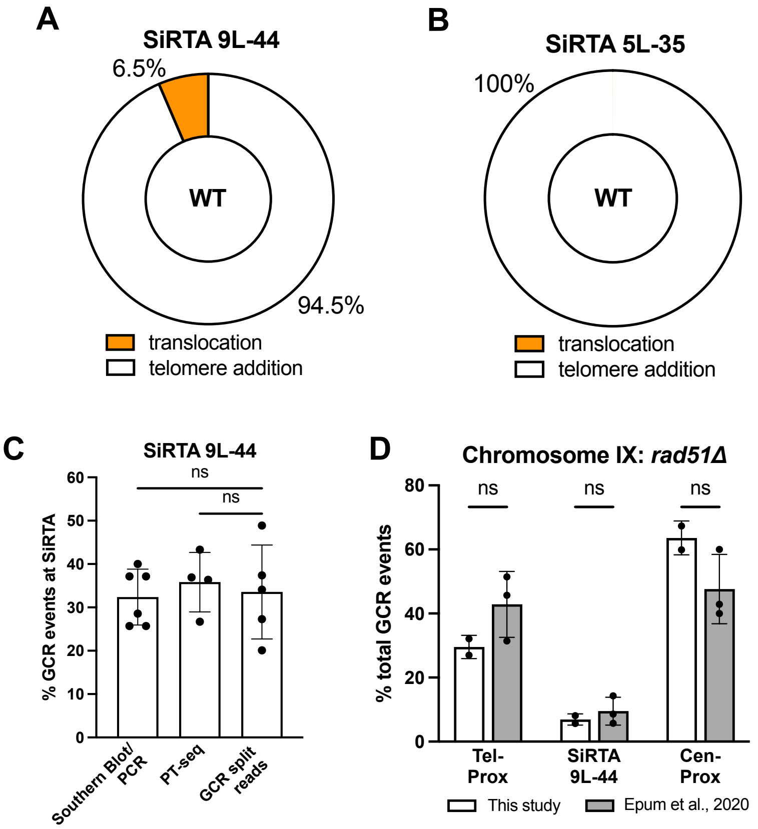


**Supplement Figure 2.** (**A**) Of the split reads that mapped to SiRTA 9L-44, the percentages of telomere addition (white) or translocations (orange) are shown. Data shown in (A) and (B) are from the same set of experiments shown in Figure 1B and 1C. (**B**) Of the split reads that map to SiRTA 5L-35, the percentages of telomere addition (white) are shown. No translocations are observed. (**C**) The percentages of GCR events at SiRTA 9L-44 are shown using three quantification methods. Each data point corresponds to 30 GCR events analyzed by Southern Blot/PCR (26), PT-seq (25), and split read analyses from this study. ANOVA with Tukey's multiple comparisons test. (**D**) The percentages of GCR events are shown for the indicated regions on chromosome IX in a *rad51Δ* strain. Quantification of GCR events by split read analysis or by Southern Blot/PCR (26) is statistically indistinguishable. Two-way ANOVA with Šídák's multiple comparisons test.

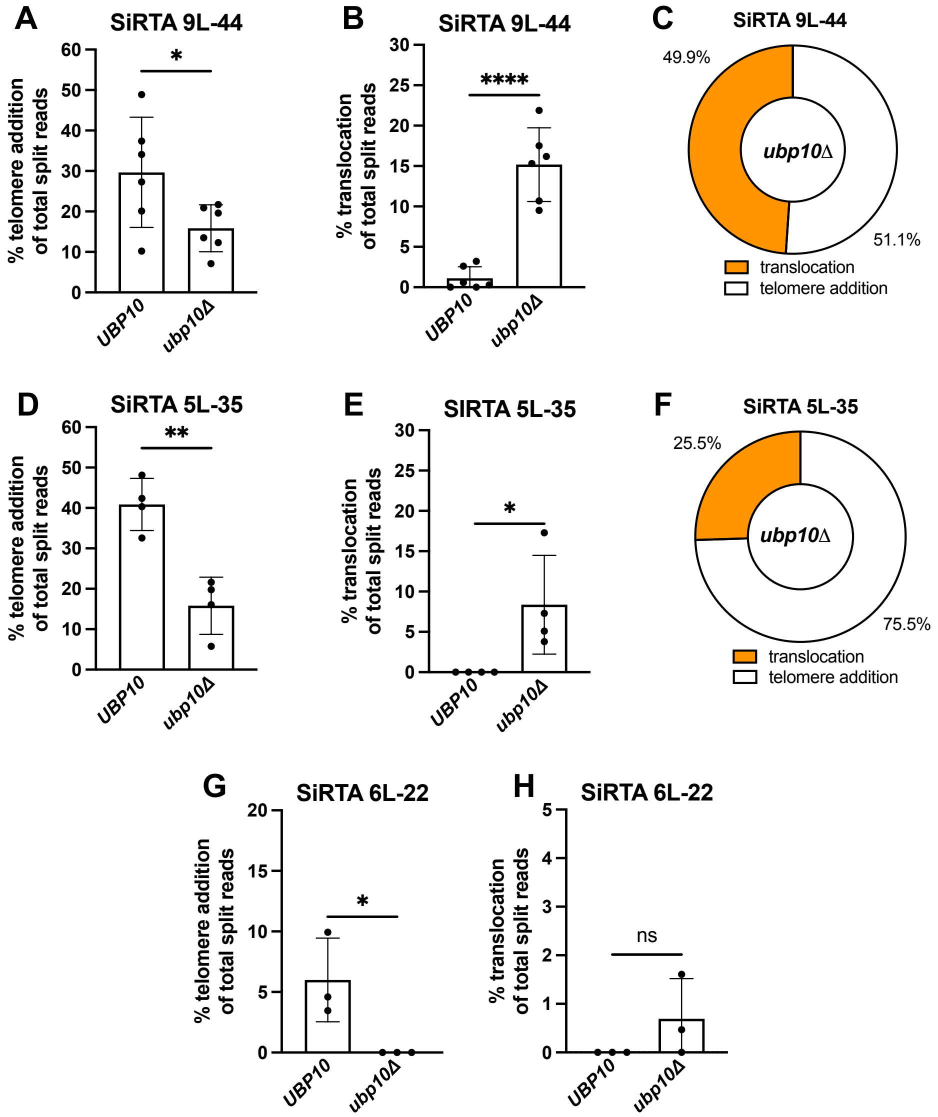


**Supplement Figure 3.** (**A-B**) Percentages of telomere addition (**A**) and translocation (**B**) split reads that map to SiRTA 9L-44 for *UBP10* and *ubp10Δ* strains are shown. Data are from 6 pools of 30 GCR events. Data is repeated from Figure 2A. * $p<0.05$, ****$p<0.0001$, t-test. (**C**) Of the split reads that map to SiRTA 9L-44, the percentages of telomere addition (white) or translocations (orange) are shown for the *ubp10Δ* strain. Data are from the same experiments shown in Figure 2A. (**D-E**) Percentages of telomere addition (**D**) and translocation (**E**) split reads that map to SiRTA 5L-35 for *UBP10* and *ubp10Δ* strains are shown. Data are from 4 pools of 40 GCR events. Data are from same experiments in Figure 2C. **$p<0.005$, *$p<0.05$, t-test. (**F**) Of the split reads that map to SiRTA 5L-35, the percentages of telomere addition (white) or translocations (orange) are shown for the *ubp10Δ* strain. Data are from the same experiments shown in Figure 2C. (**G-H**) Percentages of telomere addition (**G**) and translocation (**H**) split reads that map to SiRTA 6L-22 for *UBP10* and *ubp10Δ* strains are shown. Data are from 3 pools of 30 GCR events. Data are from same experiments in Figure 2D. *$p<0.05$, t-test.

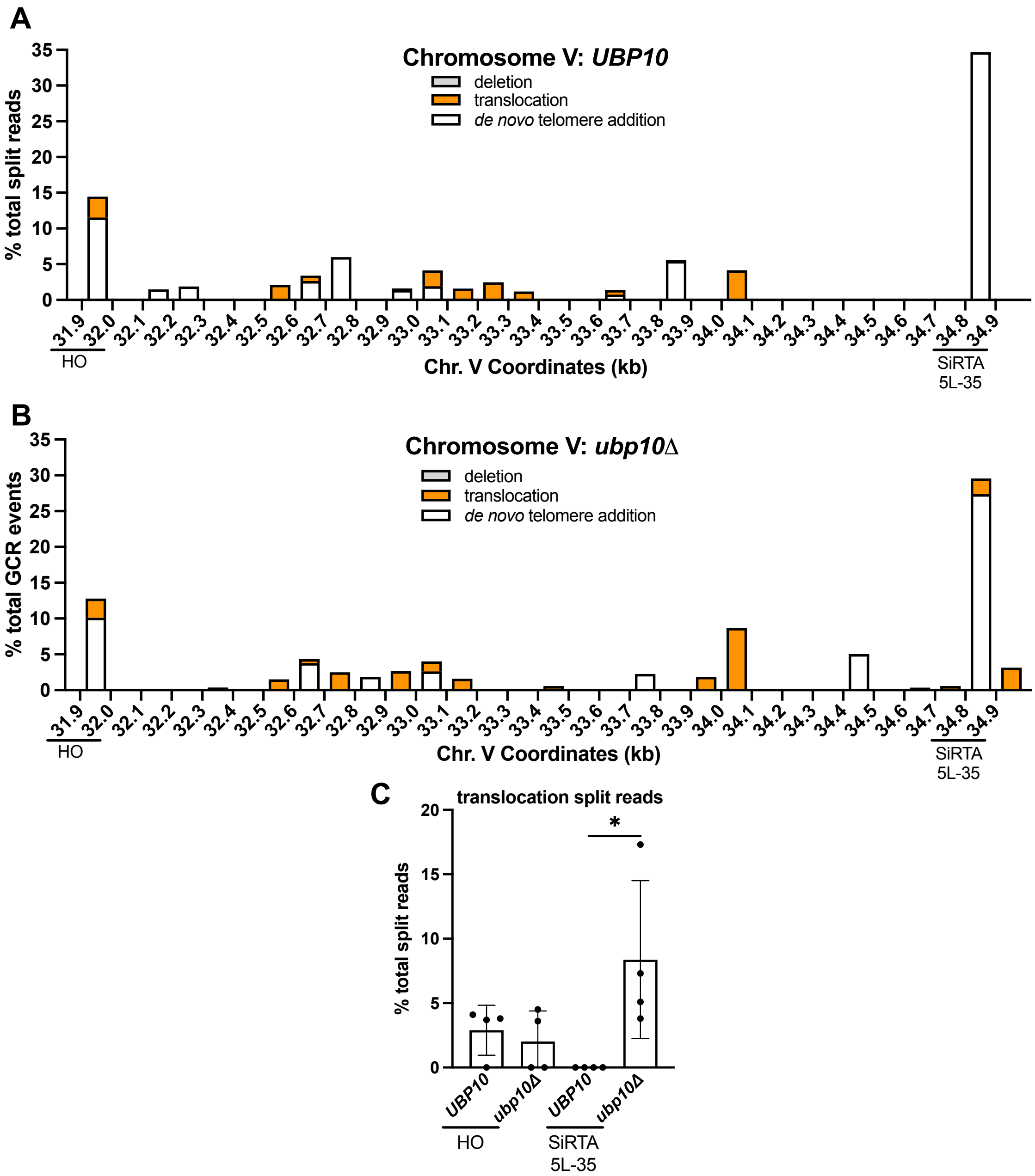


**Supplement Figure 4.** (**A-B**) Percentages of total split reads that map to each 100 bp interval between the HO cleavage site and SiRTA 5L-35 are shown. Split reads were classified as deletions (gray), translocations (orange), or telomere additions (white) as described in Methods. The X axis is the distance (in kilobases) from the left end of the chromosome. Data for *UBP10* and *ubp10Δ* strains are the same experiments described in Figure 1D and E. (**C**) Translocation split reads that map within the 100 bp region where the HO cut site was integrated (31.9-32.0) or to SiRTA 5L-35 are quantified (Supplement T3). Percentages are calculated relative to total split reads mapping from the HO site to the last essential gene. Each data point corresponds to one of the four pools of 30 GCR events as described in Figure 1D. *$p$<0.05, ANOVA with Tukey's multiple comparisons test.

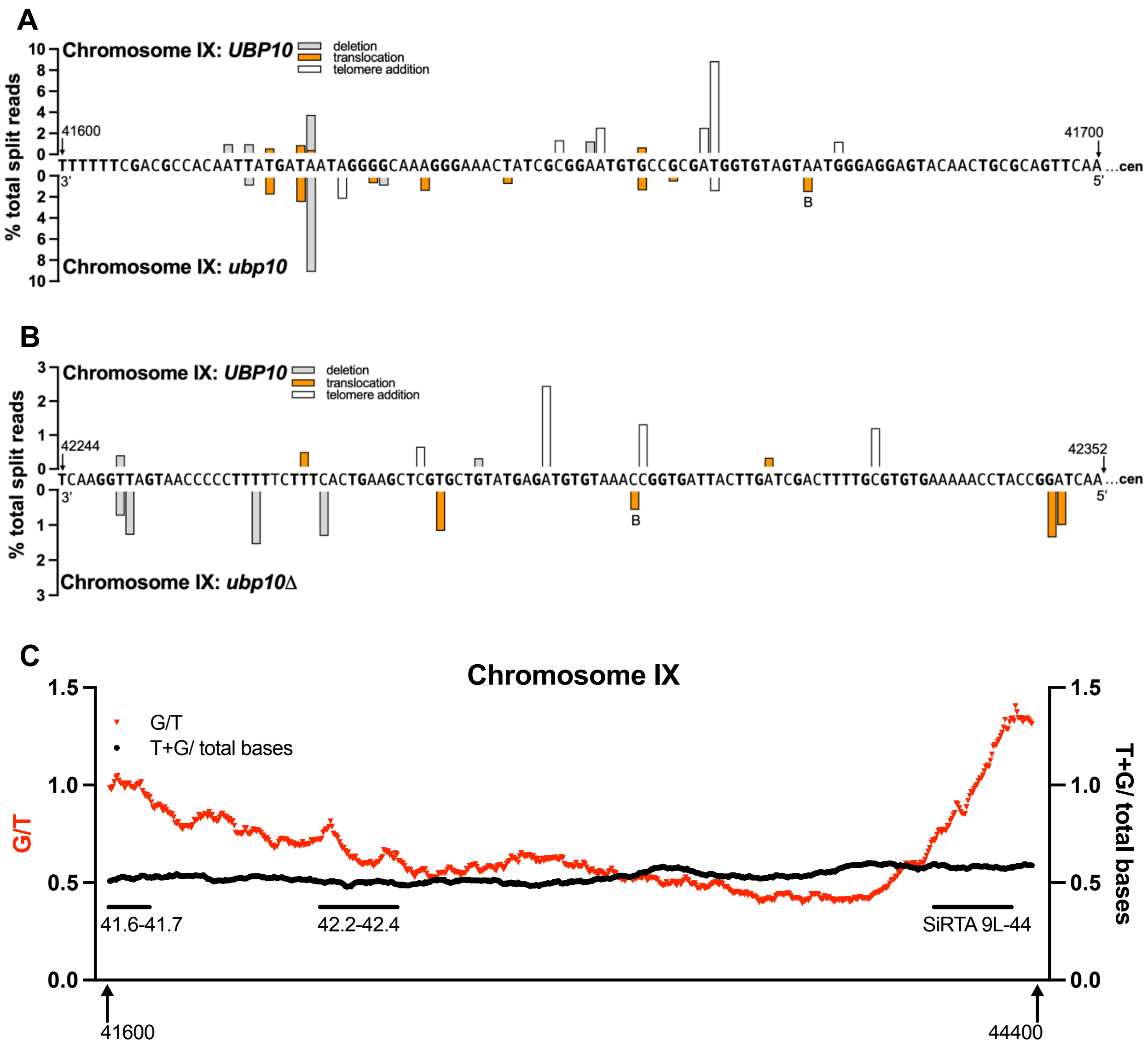


**Supplement Figure 5.** (**A-B**) Nucleotide resolution maps showing sites of telomere addition (white), translocations (orange), and deletions (gray) at the regions indicated by an arrow (A) or bracket (B) in Figure 3. Sequences are oriented 3' to 5' with the centromere located to the right (nucleotide coordinates indicated). Bars above the sequence represent data from the *UBP10* strain, bars below the sequence represent data from the *ubp10Δ* strain. (**A**) represents the region located 100 and 200 bp proximal to the HO cleavage site. (**B**) represents the region between 42.244 and 42.352 kb from the endogenous telomere. TG nucleotides are bolded in these regions. Class B translocations, denoted by the B, are observed at these sites. Unlabeled translocation bars denote translocation split reads classified at "other" translocations. (**C**) The ratio of G to T (G/T) and the fraction of T and G nucleotides (T+G/500) measured across chromosome IX 41600-44400. 500 bp intervals were analyzed as 5 nt sliding windows across the 2.8kb region beginning at the 3' end of 41600bp and extending to the 5' end of 44400bp. The ratio of thymine or guanine among nucleotides (black) and the ratio of guanine to thymine (red) are shown. Each data point corresponds to the midpoint of the 500 bp interval. The regions labelled within the graph correspond to the same regions in (**A**) and (**B**).

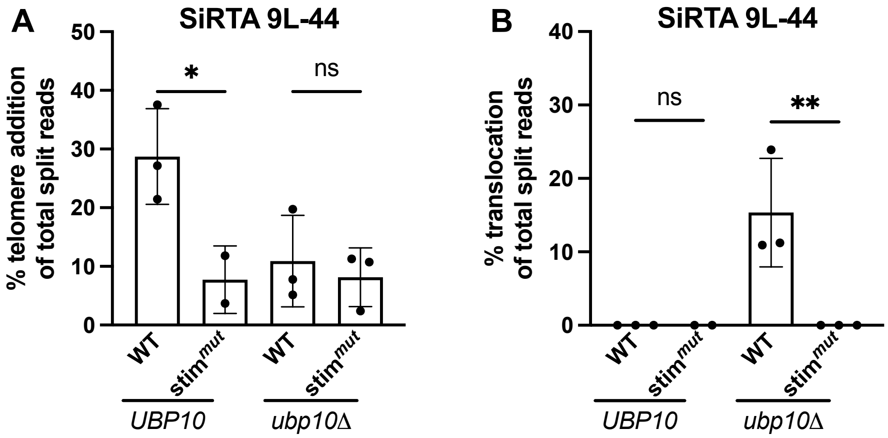


**Supplement Figure 6.** (**A-B**) Percentages of telomere addition (**A**) and translocation (**B**) split reads that map to WT SiRTA 9L-44 and SiRTA 9L-44 Stim$^{mut}$ for *UBP10* and *ubp10Δ* strains are shown. Data points represent a pooled sample of 30 GCR events. Data is repeated from experiments in Figure 4C. * *p*<0.05, ***p*<0.01, ANOVA with Tukey's multiple comparison test.

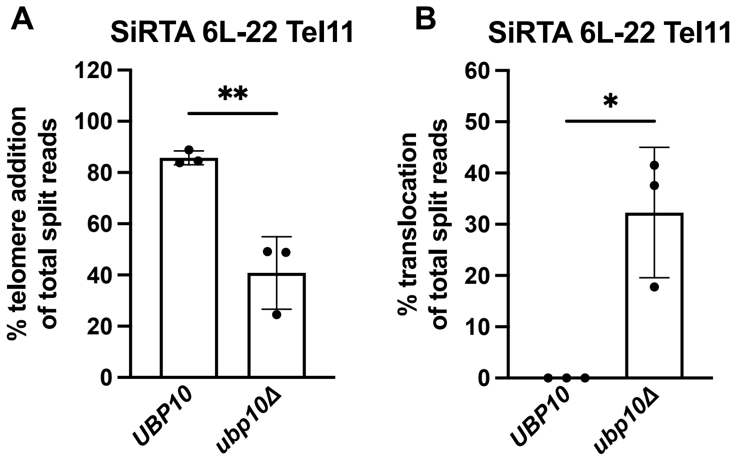


**Supplement Figure 7.** (**A-B**) Percentages of telomere addition (**A**) and translocation (**B**) split reads that map to SiRTA 6L-22 Tel11 for *UBP10* and *ubp10Δ* strains are shown. Data represent 3 pooled sample of 30 GCR events. Data is repeated from experiments in Figure 5A. ** $p<0.01$, *$p<0.05$, ANOVA with Tukey's multiple comparison test.

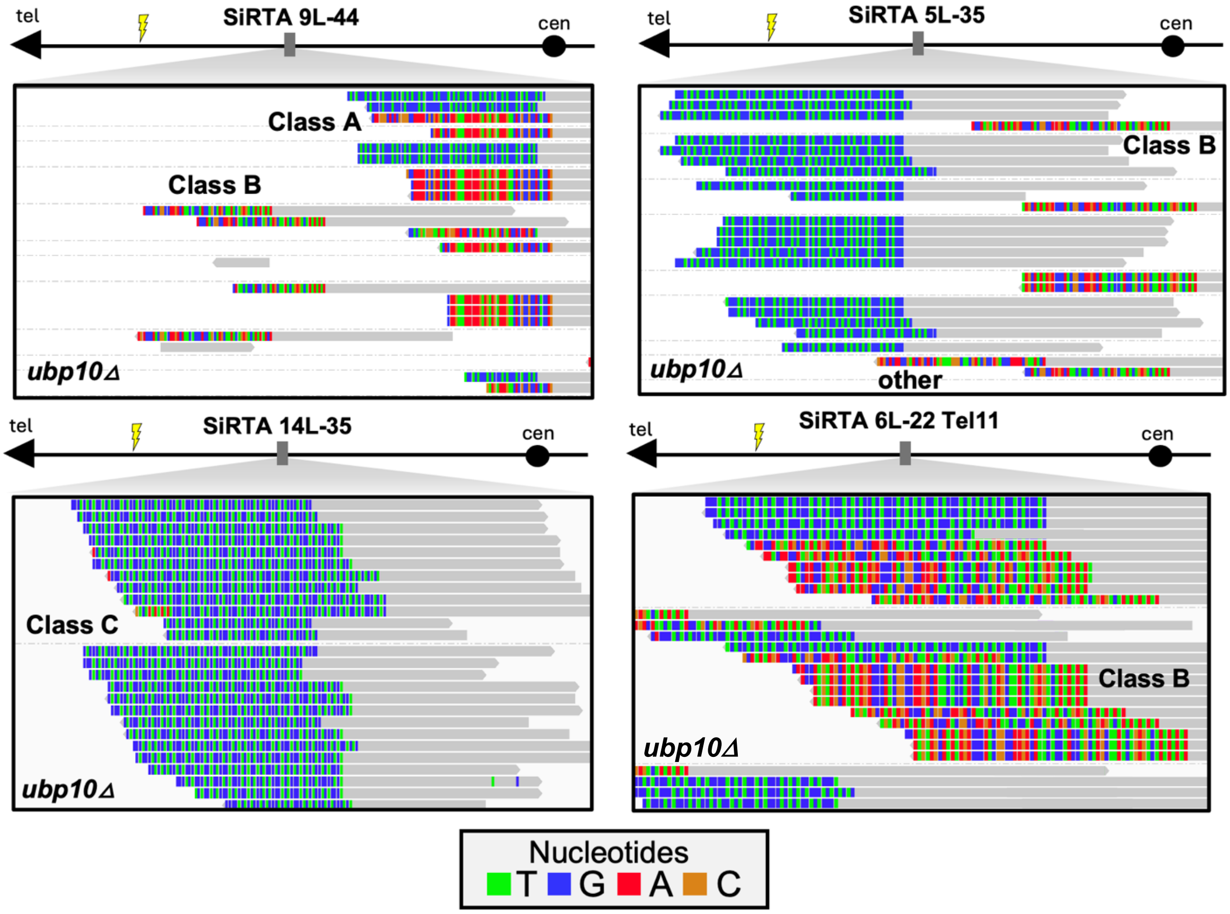


**Supplement Figure 8.** Representative images of split reads that map to SiRTAs 9L-44, 5L-35, 14L-35, and 6L-22 Tel11 in the *ubp10Δ* strain. Images are oriented with the centromere located to the right. Nucleotides in the proximal portion of the split read (gray) map accurately to the SiRTA sequence, nucleotides in the distal portion of the split read (color) align elsewhere in the reference genome. The type of translocations observed at the SiRTA are denoted in the image (see Figure 6).

**A**

**Representative Class A translocation split reads:**

**Class A split reads to Y' element:**

```
cen…CTTTGTTCATCGACAAGGACTCGGGTAAAGGCGGTGGCGGTGGTGGCAGGTGAGGCGGCGAGTTCAAATCATCATCAAAAAATTTCAAGAGTCACGCGAAAAAGGCCCCGA…tel
cen…CTTTGTTCATCGACAAGGACTCGGGTAAAGGCGGTGGCGGTGGTGGTGGTGGAACGGCG-AGTTCAAATCATCATCAAAAAATTTCAAGAGTAACGCGAAAAAG…tel
cen…CTTTGTTCATCGACAAGGACTCGGGTAAAGGCGGTGGCGGTGGTGGTGGGTGAGGCGGCGAGTTCAAATCATCATCAAAAAATTTCAAGAGTAACGCGAAAAAGGCCCCGA…tel
cen…CTTTGTTCATCGACAAGGACTCGGGTAAAGGCGGTGGCGGTGGTGGTGGTG-AGGCGGCGAGTTCAAATCATCATCAAAAAATTTCAAGAGTAACGCGAAAAAGGCCCCG…tel
cen…CTTTGTTCATCGACAAGGACTCGGGTAAAGGCGGTGGCGGTGGTGGTGGTGGAACG-ACGAGTTCAAATCATCATCAAAAAATTTCAAGAGTAACGCGAAAAAGGCCCCG…tel

cen…---CTTTGTTCATCGACAAGGACTCGGAATGCGGAAGACGTAGTGGCAGGTGAGGCGGCGAGTTCAAATCATCATCAAAAAATTTCAAGAGTAACG…tel
```

**Microhomology between recipient and donor chromosomes for Class A split reads to Y' element:**

```
cen…CTTTGTTCATCGACAAGGACTCGGGTAAAGGCGGTGGCGGTGGTGGTGGTGGAACGGCGCCACAAGTTTTAGTTTTAAATAGATCTGGAAGGG…
   …TACCATTGTTCATTCTCTAGTACTCGGAATGCGGAAGACGTAGTGGCAGGTGAGGCGGCGAGTTCAAATCATCATCAAAAAATTTCAAGAGTCACGCGAAAAAGGCCCCGA…tel

cen…CTTTGTTCATCGACAAGGACTCGGGTAAAGGCGGTGGCGGTGGTGGT-GGTGGAACGGCGCCACAAGTTTTAGTTTTAAATAGATCTGGAAGGG
   …TACCATTGTTCATTCTCTAGTACTCGGAATGCGGAAGACGTAGTGGCAGGTGAGGCGGCGAGTTCAAATCATCATCAAAAAATTTCAAGAGTAACGCGAAAAAG…tel

cen…CTTTGTTCATCGACAAGGACTCGGGTAAAGGCGGTGGCGGTGGTGGTGGTGGAACGGCGCCACAAGTTTTAGTTTTAAATAGATCTGGAAGGG…
   …TACCATTGTTCATTCTCTAGTACTCGGAATGCGGAAGACGTAGTGGCAGGTGAGGCGGCGAGTTCAAATCATCATCAAAAAATTTCAAGAGTAACGCGAAAAAGGCCCCGA…tel

cen…CTTTGTTCATCGACAAGGACTCGGGTAAAGGCGGTGGCGGTGGTGGT-GGTGGTGGAACGGCGCCACAAGTTTTAGTTTTAAATAGATCTGGAAGGG…
   …TACCATTGTTCATTCTCTAGTACTCGGAATGCGGAAGACGTAGTGGCAGGTGAGGCGGCGAGTTCAAATCATCATCAAAAAATTTCAAGAGTAACGCGAAAAAGGCCCCG…tel

cen…CTTTGTTCATCGACAAGGACTCGGGTAAAGGCGGTGGCGGTGGTGGT-GGTGGAACGCGGCGCCACAAGTTTTAGTTTTAAATAGATCTGGAAGGG…
   …TACCATTGTTCATTCTCTAGTACTCGGAATGCGGAAGACGTAGTGGCAGGTGAGGCGACGAGTTCAAATCATCATCAAAAAATTTCAAGAGTAACGCGAAAAAGGCCCCG…tel

cen…---CTTTGTTCATCGACAAGGACTCGGGTAAAGGCGGTGGCGGTGGTGGTGGTGGAACGCGGCGCCACAAGTTTTAGTTTTAAATAGATCTGGAAGGG…
   …TACCATTGTTCATTCTCTAGTACTCGGAATGCGGAAGACGTAGTGGCAGGTGAGGCGGCGAGTTCAAATCATCATCAAAAAATTTCAAGAGTAACG…tel
```

**Class A split reads to X element:**

```
cen…CATCGACAAGGACTCGGGTAAAGGCGGTGGCGGTGGTGGTGGGATTAGAGTGGTAGGGTAAGTATGTGTGTATTATTTACGATCATTTGTTAACATTTCAA…tel
```

**Microhomology between recipient and donor chromosomes for Class A split reads to X element:**

```
CATCGACAAGGACTCGGGTAAAGGCGGTGGC-GGTGGTGGTGGTGGTGGTGGAACGCGGCGCCACAAGTTTTAGTTTTAAATAGATCTGGAAGGG…
ATATCAGTATACAAGTGAGGGTGAGTGTGGCATGTGGTGGTGGGATTAGAGTGGTAGGGTAAGTATGTGTGTATTATTTACGATCATTTGTTAACATTTCAA…tel
```

**B**

**Representative Class B translocation split reads:**

```
cen…ATGAAGACGCGGATATAATGACATTTCCTAACTTTTGGGCAAAAATTCGCTATCATATGCGAGAAC----------------------------ATATATATGTCA…tel
cen…ATGAAGACGCGGATATAATGACATTTCCTAACTTTTGGGCAAAAATTCGCTATCATATGCGAGAACCGTTTGCGGAGTTTCTCGGGACAC----ATATATATGTCACTGTA…tel
cen…ATGAAGACGCGGATATAATGACATTTCCTAACTTTTGGGCAAAAATTCGCTATCATATGCGAGAACCGTTTGCGGAGTTTCTCGGGACACTAGTATATATATGTCACTGTA…tel
cen…TTTTTGGTGTTGGTGGTAATCTTCAAGCAACTGTAACAAAAGGTAGTGGTGGTTCCTATGAATCCCTATCATTTGCATGGGGGT----------ATATAT…tel
cen…TTTTTGGTGTTGGTGGTAATCTTCAAGCAACTGTAACAAAAGGTAGTGGTGGTTCCTATGAATCCCTATCATTTGCATGGGGGTTCGGTTGTATATATAT…tel
```

**Alignment between recipient and donor chromosomes without ITS:**

```
cen…TCGCTATCATATGCGAGAACATATATATGTCACTGTATTGCATGCTGGATGGTGT…tel
    TTAGGGTAGTGTTAGGGTATATATATATGTCA (IIL)
    GTTGAGAGACAGGTTAAATCATATATATGTCA (VIIR)
    ATGTGAGAGAGTGTGTGGGTATATATATGTCA (XIIR)
    AGTTGAGAGACAGGTTCATCATATATATGTCA (XVR and XVIL)

cen…TGCGGAGTTTCTCGGGACACATATATATGTCACTGTA…tel
    TTAGGGTAGTGTTAGGGTATATATATATGTCA (IIL)
    GTTGAGAGACAGGTTAAATCATATATATGTCA (VIIR)
    ATGTGAGAGAGTGTGTGGGTATATATATGTCA (XIIR)
    AGTTGAGAGACAGGTTCATCATATATATGTCA (XVR and XVIL)

cen…GAGTTTCTCGGGACACTAGTATATATATGTCACTGTA…tel
    TTAGGGTAGTGTTAGGGTATATATATATGTCA (IIL)
    GTTGAGAGACAGGTTAAATCATATATATGTCA (VIIR)
    ATGTGAGAGAGTGTGTGGGTATATATATGTCA (XIIR)
    AGTTGAGAGACAGGTTCATCATATATATGTCA (XVR and XVIL)

cen…CCTATCATTTGCATGGGGGTATATAT…tel
    TTAGGGTAGTGTTAGGGTATATATATATGTCA (IIL)
    GTTGAGAGACAGGTTAAATCATATATATGTCA (VIIR)
    ATGTGAGAGAGTGTGTGGGTATATATATGTCA (XIIR)
    AGTTGAGAGACAGGTTCATCATATATATGTCA (XVR and XVIL)

cen…GCATGGGGGTTCGGTTGTATATATAT…tel
    TTAGGGTAGTGTTAGGGTATATATATATGTCA (IIL)
    GTTGAGAGACAGGTTAAATCATATATATGTCA (VIIR)
    ATGTGAGAGAGTGTGTGGGTATATATATGTCA (XIIR)
    AGTTGAGAGACAGGTTCATCATATATATGTCA (XVR and XVIL)
```

**C**

**Representative Class C translocation split reads: X-Y' element junctions with $GT_{1-3}$sequence**

**SiRTA 14L-35**

cen…GACTCTAGTGTGGGCGGCAAGTGTTGTTGGGTGTGGGTGTGGTGTGGGTGTGTGGGTGTGGTGTGGGTGTGTGGGTGTGGGTGTGGGTGTGGGTGTGGGTATATATATGTC…tel

**SiRTA 6L-22 Tel11**

cen…TTTTTGGTGTTGGTGGTAATCTTCAAGCAACTGTAACAAAAGGTAGTGG---TGTGGGTGTGTGGTGTGTGTGGGTGTGTGGGTATATATATGT…tel
cen…TTTTTGGTGTTGGTGGTAATCTTCAAGCAACTGTAACAAAAGGTAGTGGTGG----------------------------GTGTATATATATGT…tel

**Supplement Figure 9.** Analysis of microhomology at translocation junctions. (**A**) (Top) Representative Class A translocation split reads at SiRTA 9L-44 in the *ubp10Δ* strain. Class A translocations join sequences within SiRTA 9L-44 (black) to sequences internal to the X element or the Y' element of chr. IX or chr X (orange). (Bottom) Alignment between the recipient and donor chromosomes showing microhomology. Identical nucleotides between SiRTA 9L-44 and the donor chromosome are highlighted in gray. (**B**). (Top) Representative Class B translocations at SiRTA 6L-22 Tel11 in the *ubp10Δ* strain. Class B translocations join sequences within SiRTA 6L-22 Tel11 (black) to a subset of Y' elements that begin with the ATATATAT motif (see text). (Bottom) Potential alignments between SiRTA 6L-22 Tell11 and the five chromosome termini that contain non-telomeric sequences proximal to the ATATATAT motif (2 are identical). Alignments between the SiRTA breakpoints and each donor chromosome show no extensive microhomology. (**C**) Representative Class C split reads at SiRTA 14L-35 or 6L-22 Tel11 in the *ubp10Δ* strain. These reads appear to contain sequence from the ITS that precedes the ATATATAT motif (underlined) found in donor chromosomes.

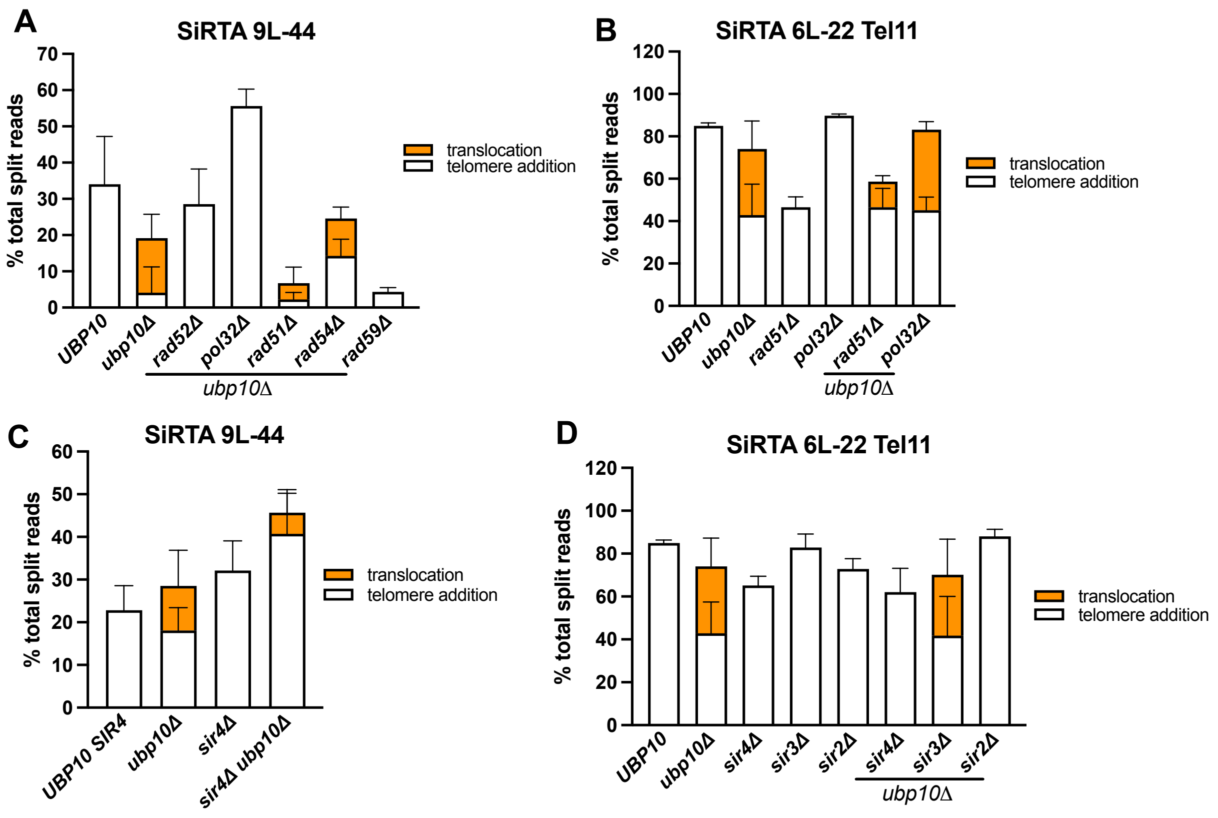


**Supplement Figure 10.** (**A&C**) Percentage of GCR split reads at SiRTA 9L-44 for each indicated strain are shown. Each bar represents the average of three experiments each containing 30 GCR events. Error bars denote SD. (**B&D**) Percentage of GCR split reads at SiRTA 6L-22 Tel11 for each indicated strain are shown. Each bar represents the average of three experiments each containing 30 GCR events. Error bars denote SD.
